# Renal Angptl4 is a key fibrogenic molecule in progressive diabetic kidney disease

**DOI:** 10.1101/2023.11.08.565844

**Authors:** Swayam Prakash Srivastava, Han Zhou, Rachel Shenoi, Myshal Morris, Leigh Goedeke, Barani Kumar Rajendran, Ocean Setia, Binod Aryal, Keizo Kanasaki, Daisuke Koya, Alan Dardik, Thomas Bell, Carlos Fernández- Hernando, Gerald I. Shulman, Julie E. Goodwin

## Abstract

Angiopoietin-like 4 (ANGPTL4) is the key protein involved in lipoprotein metabolism and has been shown to have diverse effects on tissue protection. In clinical settings, there is a reported association between higher levels of plasma Angptl4 and features of diabetic kidney disease, however, the association between kidney Angptl4 with features of diabetic kidney disease has not been well investigated. We show that both podocyte-and tubule-specific ANGPTL4 are crucial fibrogenic molecules in diabetes. Results from mRNA-array analysis in control (non-fibrotic) and diabetic (fibrotic) kidneys suggest time-dependent emergence of *Angplt4* expression. Diabetes accelerates the fibrogenic phenotype in control mice but not in ANGPTL4 mutant mice. The protective effect observed in ANGPTL4 mutant mice is correlated with a reduction in the levels of pro-inflammatory cytokines, epithelial-to-mesenchymal transition, endothelial-to-mesenchymal transition and augmented fatty acid oxidation. Mechanistically, we demonstrate that podocyte-or tubule-secreted *Angptl4* interacts with Integrin-β1 and influences the association between dipeptidyl-4 with Integrin-β1 and promotes heterodimerization of transforming growth factor-β receptor 1 (*TGFβR1*) and *TGFβR2* in cultured cells. This in turn results in *Smad3* phosphorylation and subsequent downregulation of the expression of genes involved in fatty acid oxidation; these cumulative effects led to the activation of fibrogenic phenotypes. We demonstrate the utility of a targeted pharmacologic therapy that specifically inhibits *Angptl4* gene expression in the kidneys and protects diabetic kidneys from proteinuria and fibrosis. Importantly, use of this kidney-specific targeted strategy is beneficial and does not cause any harmful effect suggesting it can be used as a novel drug molecule for treatment of diabetic kidney disease. Taken together, these data demonstrate that podocyte-and tubule-derived *Angptl4* is fibrogenic in diabetic kidneys.

## Introduction

About one-third of diabetic patients develop diabetic kidney disease, which is a leading cause of end-stage renal disease^1,2^. Kidney fibrosis is the final consequence of diabetic kidney disease^3,4^. The molecular pathways that link renal fibrosis and cardiovascular disease, two key components of diabetic nephropathy, are not well known^2,5,6^. This knowledge gap contributes to the suboptimal treatment options available for these patients. Mechanisms causing renal fibrosis include immune cell activation, pro-inflammatory cytokines, aberrant levels of chemokines, tubular cell apoptosis, endothelial cell dysfunction, podocyte cell dysfunctions, and mesenchymal activation in diverse kidney cell types^7–9^.

Though there are controversial hypotheses about myofibroblast origins, such as from epithelial cells via epithelial-to-mesenchymal transition (EMT), from endothelial cells via endothelial-to-mesenchymal transition (EndMT), from macrophages via macrophage-to-mesenchymal transition (MMT), and from resident fibrocytes from bone marrow origin, these myofibroblasts are characterized by the expression of mesenchymal markers (α-smooth muscle actin, N-cadherin and vimentin) and contribute to organ fibrogenesis ^6,9–16^. Intermediate cell types may influence mesenchymal activation processes and the health of neighboring cells^15,17–20^. Biological pathways such as activated transforming growth factor-β (TGF-β) signaling^21,22^, Notch signaling^23,24^, Wnt signaling^25–28^ and Hedgehog signaling^29–31^ lead to disruption in central metabolism and cause mesenchymal metabolic shifts in diabetic kidneys, ultimately resulting in deposition of collagens, extra-cellular matrix proteins, fibronectin, vimentin, and N-cadherin in interstitial spaces^32,33^. ANGPTL4 is known to inactivate lipoprotein lipase (LPL) activity and is expressed primarily in adipose tissues, liver, and skeletal muscle^34–36^. A major factor determining plasma triglyceride (TG) levels is endothelial-bound LPL^34,36^, which hydrolyses TG, releases non esterified fatty acids and also promotes tissue uptake of non-esterified fatty acids^34,36^. Clement et al., described two forms of ANGPTL4 : a high-isoelectric point pro-proteinuric form that is found only in glomeruli and is hyposialylated, and a sialylated, neutral isoelectric point form which is secreted from adipose tissue, liver, and skeletal muscle^37^. Conversion of the hyposialylated form to the sialylated form by treatment with N-acetyl-D-mannosamine suppresses proteinuria and the disease phenotype in a mouse model of nephrotic syndrome^37^. Studies using a podocyte-specific transgenic overexpression model that has higher podocyte-specific Angptl4 secretion demonstrated more proteinuria (∼500-fold higher), loss of glomerular basement membrane (GBM) charge and foot process effacement, whereas transgenic overexpression in adipose tissue resulted in increased circulating Angptl4, but no proteinuria, suggesting opposing effects of Angptl4 in renal tissues as compared to extra-renal tissues^37,38^. The increase in circulating Angptl4 in response to nephrotic-range proteinuria reduces the severity of the disease phenotype; however, it induces hypertriglyceridemia, suggesting a potential link between circulatory Angplt4 with proteinuria and dyslipidemia in nephrotic syndrome^38,39^. Of note, the CD-1; *db/db* mouse, which is a type 2 diabetic mouse model that mimics the renal fibrotic features of human diabetic kidney disease, has significantly higher levels of kidney *Angplt4* gene expression, suggesting a potential role of Angplt4 in diabetic kidney disease^40^.

Metabolic reprogramming which is characterized by alterations in fuel preference in different cell types is one the predominant features of mesenchymal activation in injured kidney cells^11,27,41–45^. It is still not well understood how these metabolic alterations are regulated or how they stimulate proliferation of mesenchymal cells. However, it has been shown that suppression of fatty acid oxidation (FAO) in injured epithelial cells is involved in the development of fibrosis and that abnormal glycolysis in injured endothelial cells contributes to mesenchymal activation processes^11,41,42,46,47^. Such metabolic switching is key to understanding fibrogenic programs. Genetic deficiency of Angptl4 improves glucose homeostasis and diabetes^48^ and hepatocyte-deficient mutant mice demonstrate improved hepatocyte FAO and associated improvements in obesity, diabetes, and atherosclerosis^49^. Here, our results show the crucial role of both podocyte-and tubule-specific Angptl4 in the pathogenesis of diabetic kidney disease. Targeted inhibition of kidney specific Angptl4 is a promising therapeutic option for the management of diabetic kidney disease.

## Materials and Methods

### Reagents and antibodies

Rabbit polyclonal antibody against ANGPTL4 (#40-9800) was from Invitrogen, mouse monoclonal anti-αSMA (Cat: A5228) and mouse monoclonal anti-β-actin (AC-74) (A2228) antibodies were from Sigma (St Louis, MO). Anti-TGFβR1 (Cat: ab31013), PPARα (Cat: ab215270), rabbit polyclonal anti-TGFβRII (Cat:ab61213), mouse monoclonal anti-vimentin (RV202) (Cat:ab8978), rabbit polyclonal anti-αSMA (ACTA2) (Cat:ab5694), anti-HIF1α (Cat:ab516008) and goat polyclonal anti-Snail1 (Cat:ab53519) antibodies were purchased from Abcam (Cambridge, UK). Mouse anti-β-catenin antibody (Cat:610154) was purchased from BD Biosciences. Carnitine palmitoyltransferase 1a (CPT1a) (Cat:12252), and rabbit polyclonal anti-E-cadherin antibody (24E10) (Cat:3195) antibodies were purchased from Cell Signaling Technology (Danvers, MA, USA). Anti-HSP90 was purchased from BD Biosciences (Cat:610419). Rabbit polyclonal anti-smad3 (#9513) antibody was obtained from Cell Signaling Technology (Danvers, MA, USA). Rabbit polyclonal anti-phospho-Smad3 (s423 and s425) (600-401-919) antibody was purchased from Rockland Immunochemicals (Gilbertsville, PA, USA) Fluorescence-, Alexa fluor 647-, and rhodamine-conjugated secondary antibodies were obtained from Jackson ImmunoResearch (West Grove, PA). *TGF*β*1*, *TGF*β*2*, and *TGF*β neutralizing antibodies were purchased from PeproTech (Rocky Hill, NJ).

### Animal Experimentation

All experiments were performed according to a protocol approved by the Institutional Animal Care and Use Committee at Yale University School of Medicine and Kanazawa Medical University Japan. The experiments at Yale University were in accordance with the National Institute of Health (NIH) Guidelines for the Care of Laboratory Animals and experiments at Kanazawa Medical University were carried out in accordance with Kanazawa Medical University animal protocols (#2014-89; #2013-114 and #2014-101) and approved by the Institutional Animal Care and Use Committee (IACUC). For the cell specific loss-of-function of Angptl4, we bred the Angptl4 flox/flox mice with podocin-cre mice (pmut) and Pax^rtTa^-Tet-O-Cre mice (tmut) to generate mice with deletion of Angptl4 in podocytes and tubules, respectively. All mice were on the C57BL/6 background. Six-week-old tmut mice were treated with doxycycline (2mg/ml + 3% sucrose in drinking water) for 2 weeks prior to experiments. Induction of diabetes in CD-1 mice and C57BL/6 mice was performed according to previously established experimental protocols^11,50^. A single IP dose of 200 mg/kg STZ was used to induce diabetes in CD-1 mice. In mice on the C57BL/6 background, diabetes was induced in 10-week-old male mice with five consecutive intraperitoneal (IP) doses of streptozotocin (STZ) 50 mg/kg in 10 mmol/L citrate buffer (pH 4.5). DPP-4 inhibitor (Linagliptin) was provided to 16 week-STZ-treated diabetic mice using a dose of 5 mg/kg in drinking water for 8 weeks^51^. In other experiments, control IgG and N-Integrinβ1 IgG were injected intraperitoneally to 16 week-STZ-treated diabetic mice using a dose of 1 mg/kg for 8 weeks.

In one experiment, male mice were randomized to one of four groups sixteen weeks after induction of diabetes: (i) untreated, (ii) angiotensin-converting enzyme inhibitor (imidapril-5 mg/kg in drinking water), (iii) Dichloroacetate (1g/L in drinking water), and (iv) 2-deoxyglucose 500 µg/kg via intraperitoneal injection twice per week. Mice were treated for 8 weeks and compared to untreated diabetic mice. In another set of experiments male diabetic mice were randomized to one of three groups: (i) untreated, (ii) fenofibrate (100 mg/kg by oral gavage, and (iii) simvastatin (40 mg/kg by oral gavage). All mice had free access to food and water during experiments. Blood was obtained by retro-orbital puncture. Blood glucose concentration was measured using glucose-strips. Urine albumin levels were assayed using a Mouse Albumin ELISA Kit (Exocell, Philadelphia, PA). Tissue and blood were harvested at the time of sacrifice. Some kidneys were minced and stored at −80°C for gene expression and protein analysis. Other kidneys were placed immediately in optimal cutting temperature (OCT) compound for frozen sections or 4% paraformaldehyde for histologic staining.

### Kidney specific antisense-oligonucleotide (ASO) treatment

For kidney-specific Angptl4 ASO studies, mice at 10 weeks of age were injected subcutaneously with control ASO or Angptl4 ASO at a dose of 10 or 30 mg/kg/wk for 8 weeks. Uniform chimeric 16-mer phosphonothioate oligonucleotides containing 2’,4’-constrained ethyl-D-ribose (cEt) groups at positions 1 to 3 and 14 to 16 targeted to murine Angptl4 and a control ASO were synthesized and purified by Ionis Pharmaceuticals as previously described^52^. The ASO sequences were as follows: *Angptl4* ASO: 5’-<u>AGC</u>TGTAGCAGCC<u>CGT</u> -3’ and control ASO: 5’-<u>ACG</u>ATAACGGTCA<u>GTA</u>, with the underlined sequences indicating the cEt modified bases. ASOs were dissolved in PBS for experiments.

### Blood pressure measurement

Blood pressure measurements were taken using the tail cuff method according to the manufacturer’s instructions. Briefly, mice were trained for 5 days before measurement of blood pressure. After mice were placed on the restraint platform, which was maintained at 33–34°C, the tail was placed through the optical sensor and the cuff compressed. Data are presented as the average of 10 blood pressure measurement cycles.

### Morphological Evaluation

The glomerular surface area was measured in 10 glomeruli per mouse using ImageJ software. We analyzed PAS-stained glomeruli from each mouse using a digital microscope screen. Masson’s trichrome-stained images were evaluated by ImageJ software, and the fibrotic areas were estimated^53^.

### Sirius red staining

Deparaffinized sections were incubated with picrosirius red solution for 1 hour at room temperature. The slides were washed twice with acetic acid solution for 30 seconds per wash. The slides were then dehydrated in absolute alcohol three times, cleared in xylene, and mounted with a synthetic resin. Sirius red staining was analyzed using ImageJ software, and fibrotic areas were quantified.

### Immunohistochemistry

Paraffin-embedded kidney sections (5 μm thick) were deparaffinized and rehydrated (2_min in xylene, four times; 1_min in 100% ethanol, twice; 1 min in 95% ethanol; 45_s in 70% ethanol; and 1_min in distilled water), and the antigen was retrieved in a 10_mM citrate buffer pH 6 at 98_°C for 60_min. To block endogenous peroxidase, all sections were incubated in 0.3% hydrogen peroxide for 10_min. Immunohistochemistry was performed using the Vectastain ABC Kit (Vector Laboratories, Burlingame, CA and Abcam ab64238). Rabbit polyclonal CPT1a (Abnova; H00001374-DO1P; 1:100) antibody was purchased from Abnova, USA. Rabbit polyclonal PKM2 (Cell Signaling Technology Cat# 4053, RRID:AB_1904096; 1:100), PDK4 (Abcam Cat# ab71240, RRID:AB_1269709; 1:100), PGC1α (Cell Signaling Technology Cat# 2178, RRID:AB_823600; 1:100), and CPT1a (Cell Signaling Technology Cat# 12252, RRID:AB_2797857; 1:100) antibodies were purchased from Cell Signaling Technology. Goat polyclonal anti-SIRT3 antibody (Santa Cruz Biotechnology Cat# sc-365175, RRID: AB_10710522; 1:200) was purchased from Santa Cruz Biotechnology. A goat polyclonal anti-Snail antibody (Abcam Cat# ab53519, RRID: AB_881666; 1:100) was purchased from Abcam (Cambridge, MA, USA). In negative controls, the primary antibody was omitted and replaced with the blocking solution.

### Immunofluorescence

Frozen kidney sections (5 μm) were used for immunofluorescence; double positive labeling with CD31/αSMA and E-cadherin/αSMA was measured. Briefly, frozen sections were dried and placed in acetone: methanol (1:1) solution for 10 min at − 30 °C. Once the sections were dried, they were washed twice in phosphate-buffered saline (PBS) for 5 min and then blocked in 2% bovine serum albumin/PBS for 30 min at room temperature. Then, sections were incubated in primary antibody (1:100) for 1 h and washed in PBS (5 min) three times. Next, the sections were incubated with secondary antibodies for 30 min, washed with PBS three times (5 min each), and mounted with mounting medium with DAPI (Vector Laboratories, Burlingame, CA). The stained sections were analyzed by fluorescence microscopy. For each mouse, original magnification of 400X was obtained from six different areas and quantified.

### Multiplex staining

Multiplex staining was analyzed according to the manufacturer’s instructions by using an Opal in situ kit (Waltham, MA, USA). Deparaffinized sections were labeled with E-cadherin (Opal 520 [TSA-FITC]) and Vimentin (Opal 670 [TSA-Cy5]) antibodies for EMT transition analysis while α-SMA (opal 520 [TSA-FITC]) and CD31 (opal 670 [TSA-Cy5]) antibodies were used for EndMT analysis. The cell nuclei were labeled with DAPI. In negative controls, the primary antibody was omitted and replaced with blocking solution.

### Proximity ligation Duolink in situ assay

Duolink In Situ kits were used to detect the proximity of DPP-4/integrin β1 and TGF-βR1/R2, as previously described^54,55^. Briefly, cells were passaged into 8-well culture slides (BD Biosciences) in growth medium. The cells were washed with PBS, fixed with 4% paraformaldehyde, and permeabilized with 0.2% Triton-X100. Then, blocking solution was used for 30 min at 37 °C, after which the cells were incubated in primary antibody (goat anti-integrin β1 (1:100) and rabbit anti-Angptl4 (1:100), or rabbit anti-TGFβR1 (1:500) and goat anti-TGFβR2 (1:500), or goat polyclonal DPP-4 (1:100) and rabbit anti-integrin β1 (1:100)) at 4 °C overnight. Cells were treated with two PLA probes for 1 h at 37°C, Ligation-ligase solution was added for 30 min at 37 °C, and amplification-polymerase solution was added for 100 min at 37 °C. The slides were then mounted with DAPI and analyzed by fluorescence microscopy. For each slide, original magnification of 400X was obtained from 6 different areas and quantified.

### EndMT and EMT detection

Frozen sections (5 µm) were used for the detection of EndMT and EMT. Cells undergoing EndMT were detected by double-positive labeling for CD31 and αSMA. Cells undergoing EMT were detected by double-positive labeling for E-cadherin and αSMA. Sections were analyzed and quantified by fluorescence microscopy.

### Transmission electron microscopy

For electron microscopy studies, mice were anesthetized, perfused with 4% PFA and kidneys were isolated. Kidney samples (1 mm^3^) were fixed with cacodylic acid buffer containing 1 M cacodylic acid and 25% glutaraldehyde in PBS. After Epon-embedding, an RMC/MTX ultramicrotome (Elexience) was used to cut the tissues into ultrathin sections (60–80 nm). Sections were mounted and imaged using a Nikon TE 2000U electron microscope on copper grids and stained with lead citrate and 8% uranyl acetate. A Jeol 1200 EX transmission electron microscope (Jeol LTD) equipped with a MegaView II high-resolution transmission electron microscopy camera was used to observe the copper grids. EM was performed by the Cellular and Molecular Physiology Core at Yale University. Image J was used for quantitative analysis of electron micrographs. Foot processes were measured from at least 50 μm of GBM for each mouse. Podocyte foot processes and GBM thickness were analyzed by ImageJ. Images were blinded by assigning integer numbers prior to evaluation by someone other than the scorer.

### Isolation of endothelial cells

Endothelial cells from the kidneys of non-diabetic and diabetic mice were isolated using CD31 magnetic beads. Briefly, kidneys were isolated and minced into small pieces. Using a series of enzymatic reactions by treating the tissue with trypsin and Collagenase type I solution, a single cell suspension was created. The pellet was dissolved with CD31 magnetic beads, and the CD31-labelled cells were separated with a magnetic separator. The cells were further purified on a column. Cell number was counted by hemocytometer and cells were plated on 0.1% gelatin coated Petri dishes. Cell purity was measured by flow cytometry (BD FACSDiva) using PE-conjugated CD31 (BDB553373) and FITC-conjugated CD45 (BDB553079), both from Becton Dickinson (USA)^27^.

### Isolation of kidney tubular epithelial cells (TECs)

After sacrifice, kidneys from diabetic Angptl4^-/-^ and control littermates were excised and perfused with 4% paraformaldehyde (10 mL) followed by collagenase type II digestion (2 mg/mL). After digestion, the cortical regions of the kidneys were used for further processing. The cortical region was minced and digested in collagenase buffer for an additional 5 minutes at 37°C with rotation to release cells. Digested tissue and cell suspension were passed through a 70-μm cell strainer, centrifuged at 50 g for 5 min, and washed in PBS for 2 rounds to collect TECs. Isolated TECs were seeded onto collagen-coated Petri dishes and cultured in renal epithelial cell medium (C-26130, PromoCell) supplemented with growth factors for TEC growth^27^.

### Isolation of Primary Podocytes

Podocyte isolation was performed as previously described^28^. Briefly, kidneys were dissected, minced, and digested for 45 to 60 minutes in a solution of collagenase A (1 mg/mL) (Roche) and DNAse I. The resulting suspension was strained through a 100-µm strainer and washed 3 times with Hanks Balanced Salt Solution (HBSS) buffer. Then, the suspension was resuspended in 30 mL of 45% Percoll solution (GE-Healthcare BioSciences) in isotonic buffer and centrifuged at 10,000 *g* for 60 minutes at 4°C. Glomeruli were enriched in the top band after centrifugation, and this band was collected. Cells were washed 3 times with HBSS to remove the Percoll solution. The pellet was resuspended and plated on collagen type I–coated dishes in RPMI 1640 medium with 9% FBS, 100 U/mL penicillin, 100 µg/mL streptomycin, 100 mmol/L HEPES, 1 mmol/L sodium bicarbonate, and 1 mmol/L sodium pyruvate^28^.

### Seahorse Flux Analysis of Kidneys

Basal oxygen consumption rates in kidneys from non-diabetic and diabetic wild-type and Angptl4^-/-^ mice were characterized using the Seahorse Flux Analyzer (Agilent) according to the manufacturer’s instructions. Briefly, mice were fasted overnight, sacrificed and perfused with 1x PBS. Whole kidneys were rapidly dissected, washed in 1x KHB buffer and cut into ∼2 mg pieces. Following dissection, kidney pieces were snapped into the wells of a XF Islet Capture Microplate containing 500 ml XF24 Assay Media (DMEM base media containing 1 mM pyruvate, 2 mM glutamine, 5.5 mM glucose and 100 mM palmitate, pH 7.4). Kidneys were equilibrated at 37°C for 1 h in a non-CO_2_ incubator and then assayed on a Seahorse XFe24 Analyzer (Agilent) following a 12-min equilibration period. Respiration rates were measured three times using an instrument protocol of 3-minute mix, 2-minute wait, and 3-minute measure. Flux rates were normalized to tissue weight. Experiments were repeated in 3 mice per condition.

### ATP measurement

ATP content was determined using the ATP Colorimetric Assay kit (Biovision), following the manufacturer’s instructions.

### mRNA Array Analysis

Total RNA was isolated using RNeasy Mini Kit (QIAGEN, Hilden, Germany). The RNA concentration was analyzed at 260/280 nm by photometry. The sense cDNA was prepared using a kit from Ambion (Austin, TX) and target hybridizations were analyzed using a Mouse Gene 1.0 ST Array (Affymetrix, Santa Clara, CA). Hybridization was performed for 17 h at 45°C in a GeneChip Hybridization Oven 640 (Affymetrix). After washing and staining in a GeneChip Fluidics Station 450, hybridized cDNAs were detected using the GeneChip Scanner 3000. The digitalized image data were processed using the GeneChip Operating Software version 1.4. After hierarchical clustering, the results were presented as a heat map. PANTHER biological classification system was used to select specific function-related genes, pathway analysis and molecular function of selected genes.

### mRNA isolation and qPCR

Total RNA was isolated using standard Trizol protocol. RNA was reverse transcribed using the iScript cDNA Synthesis kit (Bio-Rad) and qPCR was performed on a Bio-Rad C1000 Touch thermal cycler using the resultant cDNA, as well as qPCR Master mix and gene specific primers. The list of mouse primers used is in **Table S1**. Results were quantified using the delta–delta-cycle threshold (Ct) method (ΔΔCt). All experiments were performed in triplicate and 18S was utilized as an internal control.

### RNA Extraction and microRNA Array Analysis

Frozen kidney tissues were first placed in RNA Later (Life technologies) for 16h at −20^◦^C before the subsequent homogenization process to avoid RNA degradation while extracting high-quality microRNA. Total RNA was isolated using the miRNeasy Kit (Qiagen, Hilden, Germany). The RNA was quantified with a nanodrop spectrophotometer (ND-1000, Nanodrop Technologies, Wilmington, DE, USA). Ratios of OD 260/280 were between 1.9 and 2.0. The integrity of RNA samples was determined usimg a Bioanalyzer 2100 (Agilent, Santa Clara, CA, USA) and all samples gave RIN values in the range of 8.0–8.5. The input for the Agilent microRNA labeling system was 100 ng total RNA. Dephosphorylated and denatured total RNA was labeled with cyanine 3-pCp and subsequently hybridized to the Agilent Mouse microRNA Microarray Release 15.0 using the microRNA Complete Labeling and Hyb Kit (Agilent Technologies, Inc., Santa Clara CA). Following hybridization for 20 h, the slides were washed with Gene Expression Wash Buffer Kit (Agilent, Santa Clara, CA, USA) and measured using an Agilent Scanner G2565BA. Agilent Feature Extraction Software 9.5.1 and GeneSpring GX software 12.5 (Agilent) were used for data processing, analysis, and monitoring. Predicted targets, as extracted from www.microRNA.org, were used to identify the potential putative targets of microRNAs^56^ for cellular pathways and annotation enrichments were identified using PANTHER databases. These databases categorize a set of genes from the input gene lists based on annotation similarity and then map them as significantly over-represented in a biological pathway, thus suggesting that the enriched pathways might play a role in the physiological condition being considered.

### microRNA RNA Isolation and qPCR

The complementary DNA was synthesized by a miScript II RT kit (Qiagen) using the hiSpec buffer method. microRNA expression was quantified using the miScript SYBR Green PCR Kit (Qiagen) using 3 ng of complementary DNA. The primers to quantify Mm_miR-29a, Mm_miR-29b, and Mm_miR-29c were the miScript primer assays pre-designed by Qiagen. The mature microRNA sequences were 5’UAGCACCAUCUGAAAUCGGUUA for Mm_miR-29a, 5’UAGCACCAUUUGAAAUCAGUGUU for Mm_miR-29b, and 5’UAGCACCAUUUGAAAUCGGUUA for Mm_miR-29c. Hs_RNU6-2_1 (Qiagen) was utilized as an internal control.

### *In vitro* experiments and transfection

Human HK-2 cells were used at passages 4–8 and cultured in epithelial basal media with growth factors and 10% serum. Human Angptl4-and Angptl3-specific siRNA (Invitrogen) were used at a concentration of 40 nM for 48 h to effectively knock down Angptl4 and Angplt3. Cells were treated with or without TGFβ2 (10 ng/ml) for 48h and harvested for western blot analysis. In a second set of experiments, isolated tubules from diabetic control and tmut mice were co-cultured with podocytes and endothelial cells. Similarly, isolated podocytes from diabetic control and pmut mice were co-cultured with tubules and endothelial cells. In another experiment, Human HK-2 cells were cultured in DMEM and Keratinocyte-SFM (1X) (Life Technologies Green Island NY) media, respectively. When the cells on the adhesion reagent reached 70% confluence, 10 ng/ml recombinant human TGFβ1 was placed in the serum-diluted medium for 48 h.

For the transfection studies, HK-2 cells, which were maintained in serum-diluted media, were passaged in 6-well plates. HK-2 cells were then transfected with 100 nM of antagomir for miR-29a, miR-29b and miR-29c (Qiagen), using Lipofectamine 2000 transfection reagent (Invitrogen, Carlsbad, CA, USA). The cells were incubated for 6 h with lipofectamine and the anti-microRNA complex in antibiotic-free media, after which the medium was refreshed before the cells were incubated for another 48 h. Upon the termination of the incubation, total RNA was isolated using the miRNeasy Kit (Qiagen) following the manufacturer’s instructions for the expression analysis, lipid uptake and lipid oxidation.

### Luciferase assay

A luciferase assay was used to analyze the activity of the 3’ untranslated region (3’UTR) in human Angptl4. Amplified vector DNA (Cloned 3’UTR human Angptl4 (Endofectin GeneCopoeia catalog no. EF013-S) and miRNA 3’UTR (MmiT088761-MT06-264ng/ul) were transfected into HK-2 cells. Transcriptional activity was evaluated by Dual-Luciferase Reporter Assay System (Promega) in triplicate samples in the presence of control microRNA, mimetic, antagomiR, or inhibitor of 29s (100nM),

### Fatty acid uptake

Cultured isolated kidney endothelial cells were incubated with medium containing 2 μCi [^14^C]-palmitate. [^14^C]-palmitate uptake was measured by liquid scintillation counting.

### Fatty Acid Oxidation

Cultured isolated kidney endothelial cells were incubated with medium containing 0.75 mmol/L palmitate (conjugated to 2% free fatty acid–free BSA/[^14^C] palmitate at 2 μCi/mL) for 2 h. One mL of the culture medium was transferred to a sealable tube, the cap of which housed a Whatman filter paper disc. ^14^CO_2_ trapped in the media was then released by acidification of media using 60% perchloric acid. Radioactivity that had become adsorbed onto the filter discs was quantified by liquid scintillation counting.

### Statistical analysis

All values are expressed as mean ± SEM and analyzed using GraphPad Prism 7 (GraphPad Software, Inc., La Jolla, CA). One-way ANOVA followed by Tukey’s test and two-way ANOVA were employed to analyze significance when comparing multiple independent groups. The post hoc tests were run only if F achieved P< 0.05 and there was no significant variance in homogeneity. In each experiment, N represents the number of separate experiments (in vitro) or the number of mice (in vivo). Technical replicates were used to ensure the reliability of single values. Data analyses were blinded. The data were considered statistically significant at p < 0.05.

## Results

### Loss of Angptl4 protects against a fibrotic phenotype in a murine model of progressive diabetic kidney disease

Streptozotocin-induced diabetic mice are the established mouse model for the studying diabetic kidney disease (DKD). Diabetes mellitus caused gain of time-dependent fibrogenic kidney phenotypes, a higher rate of EMT, as evidenced by higher vimentin expression in E-cadherin positive cells, and a higher rate of EndMT, as shown by higher αSMA in CD31-positive cells, at 24-weeks post-streptozotocin injection, when compared to nondiabetic control mice (**Figure 1A-B**). To study cellular changes more broadly in DKD, we performed mRNA array analysis in control and fibrotic diabetic kidneys. Array data demonstrate that there were 284 genes upregulated (>2 fold) and 150 genes downregulated (>0.5 fold) in diabetes and these altered genes were associated with fibrosis-related biological functions and pathways (**Figure 1C**, **Figure S1 and Figure S2)**. Interestingly, we found significant alteration in the expression of genes regulating LPL. LPL was significantly downregulated and the LPL regulators *Angptl3* and *Angplt4* were significantly up regulated in diabetes (**Figure 1C**). To test the role of LPL in the phenotype of DKD, we used a chemical inhibitor of LPL, poloxamer 407, in diabetic mice which have early onset kidney fibrosis. Poloxamer 407 treatment suppressed LPL activity and increased the level of plasma triglycerides. However, it did not change the severity of kidney fibrosis when compared to un-treated mice, suggesting that suppression in LPL activity is not related to fibrosis in diabetic kidneys **(Figure S3)**. The mRNA expression levels of the LPL regulators, *Angptl3* and *Angplt4* were higher in time-dependent fibrotic kidneys, similar to the pattern observed in fibronectin. *Angptl3* and *Angplt4* mRNA expression was found to be higher in high glucose-(30 mM) treated cultured human tubular HK-2 cells than in low-glucose conditions (**Figure 1D** **and Figure S4A)**. *Angptl4* knockdown abolished the TGFβ1-associated expression of fibrogenic genes in high glucose-stimulated cells while *Angptl3* did not have such a prominent effect, suggesting that *Angptl4* is a key contributor to fibrogenesis in tubule cells **(Figure S4B)**.

**Figure 1.**
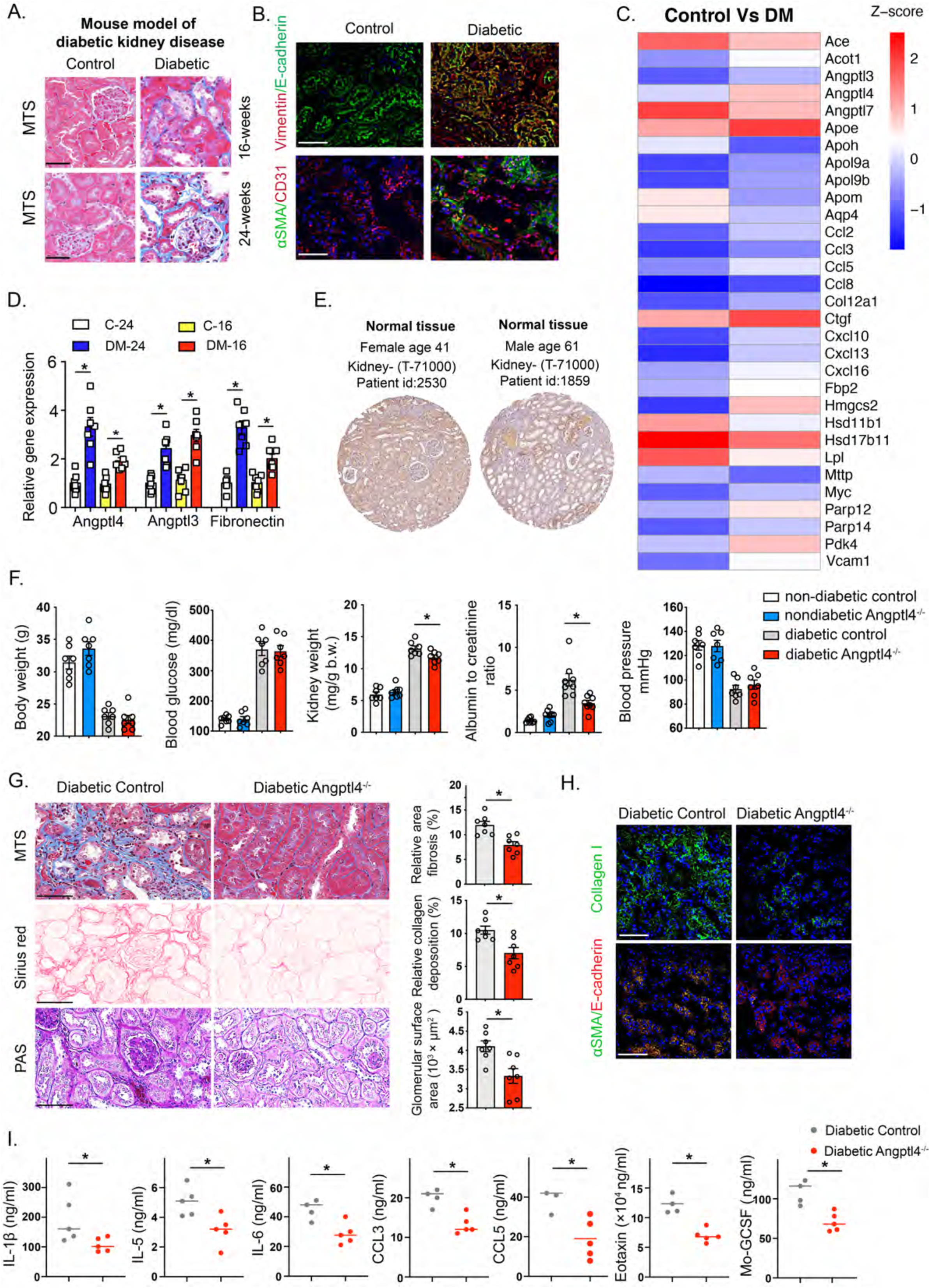
Upregulated Angptl4 expression is associated with diabetic kidney disease. **A.** Masson trichrome staining in kidneys of CD-1 mice at 16-weeks and 24-weeks post streptozotocin injection. Scale bar 50 μm. **B.** Immunofluorescence analysis of vimentin/E-cadherin and αSMA/CD31 in kidneys from nondiabetic (control) and diabetic mice. FITC-labeled E-cadherin, rhodamine-labeled vimentin and DAPI (nuclei, blue); FITC-labeled αSMA, rhodamine-labeled CD31 and DAPI (nuclei, blue), were used. Representative merged (original magnification 400X) images are shown. Scale bar 50 μm.. **C.** mRNA array analysis in control and diabetic mice. Heat map of analyzed gene expression is shown. **D.** Relative gene expression levels of *Angptl4*, *Angplt3*, and *fibronectin* in control and diabetic mice. n=7 mice/group. **E.** Immunohistochemical images of Angptl4 expression in human kidney tissue from https://www.proteinatlas.org. **F.** Physiological characteristics in nondiabetic and diabetic control and Angptl4^-/-^ mice. n=7 mice/group except for blood pressure where n=6 mice/group. Two replicate experiments were analyzed. **G.** Histologic images of Masson trichrome, Sirius red and PAS staining in diabetic control and Angptl4^-/-^ mice (original magnification 300X). Relative area of fibrosis, relative collagen deposition and glomerular surface area were calculated using ImageJ. Representative images are shown. Scale bar 100 μm. n=7 mice/group. **H.** Immunofluorescence analysis of collagen I and αSMA/CD31 was performed in kidneys from diabetic control and diabetic Angptl4^-/-^ mice. FITC-labeled collagen I, FITC-labeled αSMA, rhodamine-labeled E-cadherin and DAPI (nuclei, blue), were used. Representative merged (original magnification 200X) images are shown. Scale bar 50 μm. n=7 mice/group. Three independent experiments were analyzed. **I.** Plasma levels of IL-1β, IL-5, IL-6, CCL3, CCL5, eotaxin and Mo-GCSF from diabetic control and diabetic Angptl4^-/-^ mice were analyzed by cytokine array analysis. n=5 mice/group. Data are mean ± SEM. One-way Anova with Tukey’s multiple comparison test was used to calculate statistical significance. Significance-\**p*<0.05.

The metabolic function of *Angptl4* in regulating proteinuria and its relationship with hypertriglyceridemia in a mouse model of glomerular disease have been described in the past^37,38^. *Angptl4* is highly expressed in the human kidney **(Figure S4C)**, though its function in regulating metabolism in diabetes mellitus is poorly understood. Using human data from a publicly-available human atlas (https://www.proteinatlas.org/), it is evident that *Angptl4* is expressed in glomerular cells and tubules in both males and females (**Figure 1E**). To investigate the role of *Angplt4* in diabetic kidney disease, we used Angptl4 mutant (Angplt4^-/-^) mice. Angptl4^-/-^ mice and their littermates were born at expected Mendelian ratios. We did not observe any remarkable differences in body weight, blood glucose, kidney weight, albumin-to-creatine ratio or blood pressure in the non-diabetic mice. Blood glucose, kidney weight and albumin-to-creatine ratios were significantly higher in diabetic control mice. Blood pressure was significantly lower in diabetic mice of both genotypes, as expected (**Figure 1F**). There were no differences in fibrosis levels between the non-diabetic control and Angplt4^-/-^ mice **(Figure S5)**. Diabetes increased the area of fibrosis, relative collagen deposition and extent of glomerulosclerosis in the kidneys of control mice; however, the same effect was not observed in the kidneys of Angplt4^-/-^ mice (**Figure 1G**).

Immunofluorescence analysis revealed higher collagen I deposition in kidneys from diabetic control mice when compared to kidneys from diabetic Angplt4^-/-^ mice. Diabetes enhanced the rate of EMT in the kidneys of control mice as evidenced by increased co-localization of αSMA in E-cadherin positive cells; this effect was not prominent in kidneys from Angplt4^-/-^ mice (**Figure 1H**). Aberrations in cytokine and chemokine levels are associated with mesenchymal transformation in tubules and endothelial cells; therefore, plasma cytokines levels were measured. Angplt4^-/-^ mice had suppressed levels of proinflammatory cytokines and chemokines, such as IL-1β, IL-5, IL-6, CCL3, CCL5, eotaxin and Mo-GCSF, when compared to diabetic control mice (**Figure 1I**).

The phenotype of these Angplt4^-/-^ mice was also evaluated using an accelerated model of nephropathy, which combined the effects of both the streptozotocin-induced diabetic model as well as the UUO model^28^. In this model, control and Angplt4^-/-^ mice were subjected to UUO followed by 4 daily doses of streptozotocin and euthanasia on day 12. There were no differences in blood glucose concentrations between the two groups but UUO-operated Angplt4^-/-^ mice demonstrated lower kidney weights and lessened fibrosis when compared to UUO-operated control mice **(Figure S6A-B)**.

### Metabolic reprogramming by Angptl4 deficiency protects against diabetic kidney disease

Metabolic shifts or “switches’ play critical roles in the health and disease processes of kidney cells and recent studies suggest that such alterations in fuel preference are associated with mesenchymal transformation in kidney cells ^11,41,43,57–59^. Our data show that diabetes suppressed the expression of proteins associated with FAO such as carnitine palmitoyltransferase 1a (CPT1a), peroxisome proliferator activated receptor gamma coactivator 1-alpha (PGC1α), and sirtuin 3 (SIRT3) in the kidneys from diabetic control mice but not in the kidneys from Angplt4^-/-^ mice (**Figure 2A**). Diabetes increased the expression levels of regulators that control the oxidation of fuel types including pyruvate kinase muscle type 2 (PKM2), pyruvate dehydrogenase kinase 4 (PDK4), hypoxia inducible factor 1-alpha (HIF1α) and the EMT regulator Snail1 in the kidneys from diabetic control mice whereas these effects were not prominent in kidneys from Angplt4^-/-^ mice (**Figure 2B-C**). These data are aligned with the results of FAO and metabolic flexibility assays in these kidneys. The kidneys from diabetic control mice demonstrated a diminished level of FAO and basal oxygen consumption rate (OCR) whereas the kidneys from diabetic Angplt4^-/-^ mice demonstrated an increase in FAO and R levels **(Figure 2E; Figure S6C)**. In addition, the levels of FAO were significantly suppressed in tubules, endothelial cells, and podocytes in diabetic mice when compared to those cell types in nondiabetic mice. Tubules, endothelial cells, and podocytes from diabetic Angplt4^-/-^ mice had elevated levels of FAO when compared to diabetic control mice, suggesting a critical role of *Angptl4* in the regulation of FAO in diabetic kidneys (**Figure 2E**).

**Figure 2.**
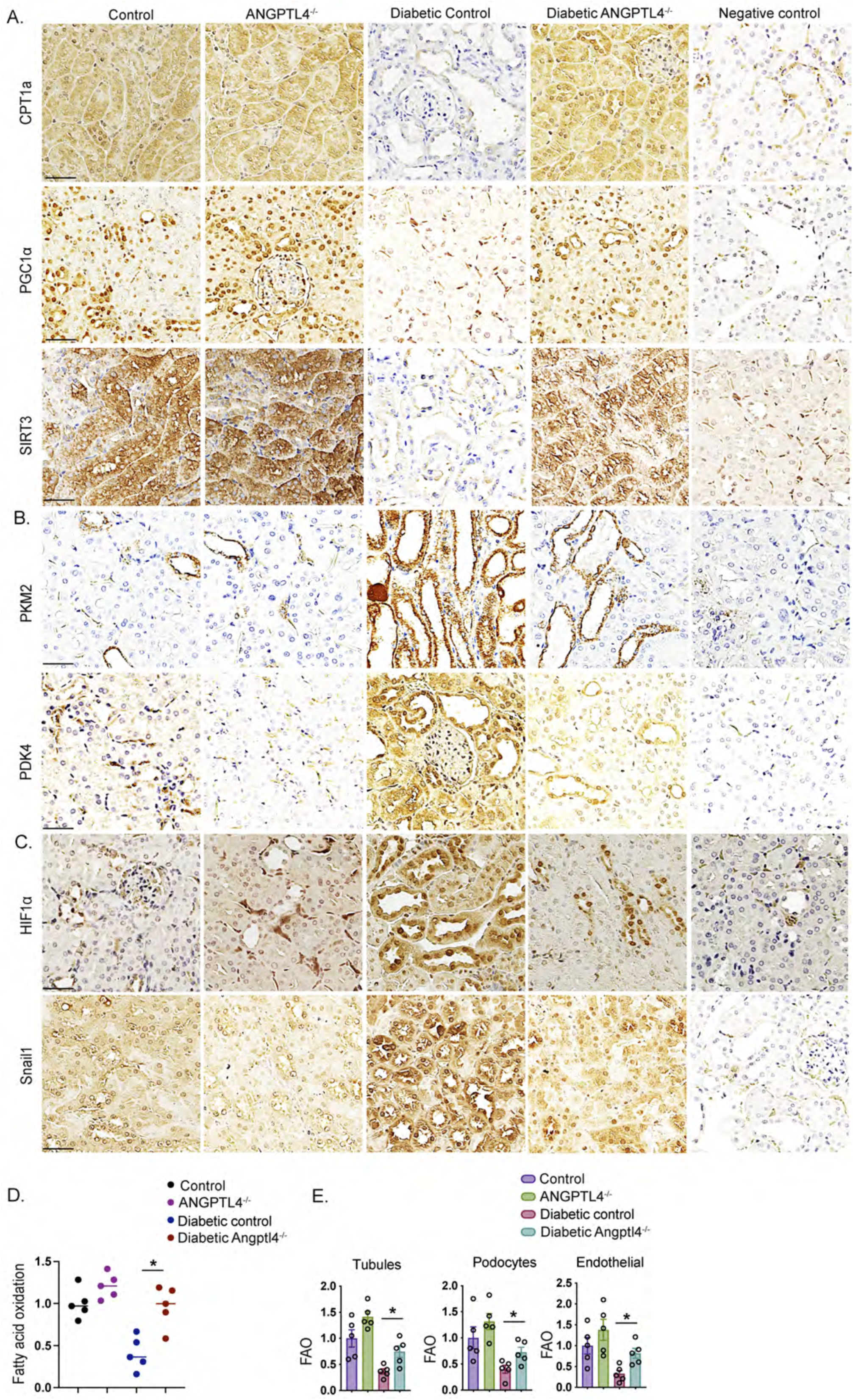
Metabolic reprogramming by Angptl4 deficiency protects against diabetic kidney disease. **A-C.** Immunohistochemical analysis of, CPT1a, PGC1α, SIRT3, PKM2, PDK4, HIF1α and Snail1 expression in kidneys from nondiabetic and diabetic, control and Angptl4^-/-^ mice. Scale bar 50 μm. n=6 mice/group. Two independent replicate experiments were performed. Representative micrographic images (original magnification 300X) are shown. **D.** Radiolabeled [^14^C]palmitate oxidation was measured and and [^14^CO_2_] release were measured. CPM of each sample was counted. n=5/group. **E.** Fatty acid oxidation analysis in isolated tubules, podocytes and endothelial cells from nondiabetic and diabetic control and Angptl4^-/-^ mice. Radiolabeled [^14^C]palmitate oxidation and [^14^CO_2_] release were measured. CPM of each sample was counted (n=5/group). Data are mean ± SEM. One-way Anova with Tukey’s multiple comparison test was used to calculate statistical significance. Significance-\**p*<0.05.

### Angptl4 deficiency in podocytes suppresses fibrogenic phenotypes

To test the role of *Angptl4* in podocytes, we generated podocyte-specific Angptl4 mutant mice (pmut) by crossing Angptl4^fl/fl^ mice with mice carrying the *podocin Cre* driver (Angptl4^fl/fl^; *podocin Cre+*) (**Figure 3A**). Pmut mice had significantly suppressed levels of *Angptl4* expression in isolated podocytes when compared to control mice **(Figure S7A)**. Nondiabetic and diabetic pmut mice did not show any remarkable alterations in body weight, blood glucose or kidney weight when compared to their nondiabetic and diabetic controls, respectively (**Figure 3B**). However, diabetic pmut mice had suppressed ACRs, relative fibrosis and glomerulosclerosis levels when compared to diabetic control mice (**Figure 3B,C**). Glomerular ultrastructure was analyzed by transmission electron microscopy and diabetic control mice displayed some podocyte foot process effacement while pmut did not show such a prominent effect (**Figure 3D**).

**Figure 3.**
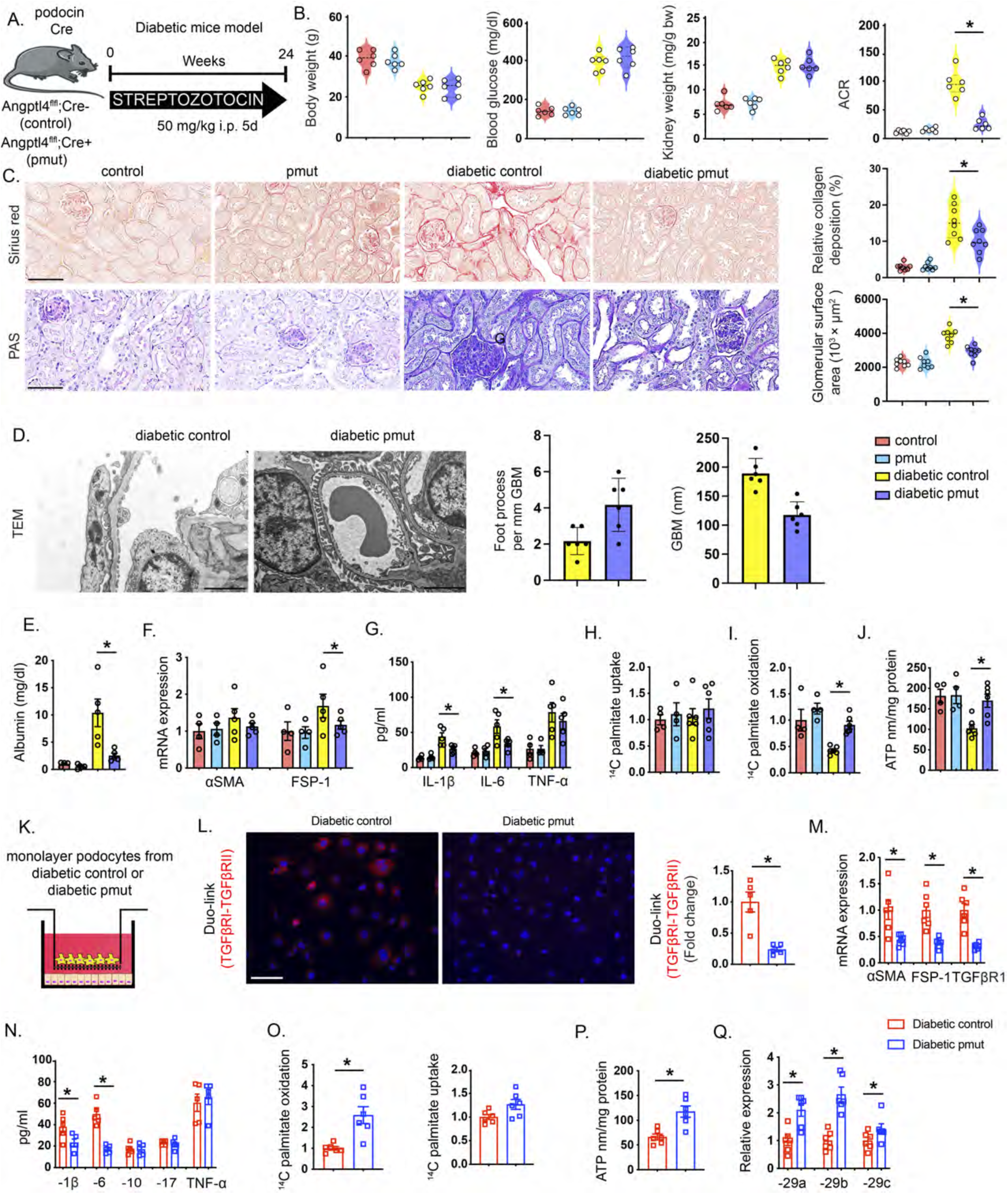
Metabolic reprogramming by podocyte Angptl4 loss protects against diabetic kidney disease. **A.** Schematic diagram, showing induction of diabetes in *Angptl4 fl/fl; Podocin Cre-* (Control) and in *Angptl4 fl/fl; podocin Cre+* (pmut) mice. Five doses of STZ (50 mg/kg/day IP) were injected to induce diabetes in mice; after 16 weeks kidneys were evaluated for fibrosis. Art was created using Servier medical art illustrations. **B.** Physiological parameters in diabetic and non-diabetic mice of both genotypes. n=6 mice/group combined from two replicate experiments were evaluated. **C.** Histologic images from Sirius red and PAS staining from kidneys of non-diabetic and diabetic control and pmut mice. Representative images (original magnification 300X) are shown. Relative area of fibrosis (%) and relative collagen (%) were measured using the ImageJ program. Scale bar 50 µm. **D.** Representative transmission electron microscopy images are shown. n=3/group. Scale bar 1 μm. Relative density of podocyte foot processes and glomerular basement membrane (GBM) thickness were calculated using ImageJ software. Six independent images of the staining were analyzed. **E.** After 48 h, albumin was measured in the cell media of cultured isolated podocytes from nondiabetic control and pmut, and diabetic control and pmut mice. **F.** qPCR mRNA gene expression of α*SMA* and *FSP-1* in isolated podocytes from nondiabetic control and pmut and diabetic control and pmut mice. n=4/non-diabetic group and n=5/diabetic group. **G.** IL-1β, IL-6, and TNFα levels measured in culture media. **H.** Radiolabeled [C^14^]palmitate uptake analysis and **I.** [^14^C] palmitate oxidation in isolated podocytes from indicated groups. CPM of each sample was counted. **J.** ATP measurement via calorimetric assay in isolated podocytes. **K.** Schematic diagram showing co-culture experimental set-up. Isolated podocytes from diabetic control and diabetic pmut mice were cocultured with isolated tubules from control mice. **L.** Representative Duolink *In Situ* images of TGF-βR1/2 from tubules that were co-cultured with podocytes either from diabetic control or diabetic pmut mice and corresponding quantification. Combination sites of TGF-βR1/2 in cells were counted in 10 different areas per condition (original magnification 400X). **M.** qPCR gene expression analysis of *FSP-1*, α*SMA* and *TGF*β*R1.* **N.** Measurement of IL-1β, IL-6, IL-10, IL-17 and TNFα levels in the cell supernatant. **O.** Radiolabeled [^14^C]palmitate uptake and oxidation, were measured in indicated groups. CPM of each sample was counted. **P.** ATP measurement in isolated tubules **Q.** qPCR gene expression analysis of miR-29a-3p, miR-29b-3p and miR-29c-3p in the tubules cocultured with podocytes from diabetic control and diabetic pmut mice. Data are mean ± SEM. One-way Anova with Tukey’s multiple comparison test was used to calculate statistical significance. Significance-\**p*<0.05.

Podocytes from diabetic pmut mice had suppressed albumin permeability when compared to podocytes from diabetic control mice as measured by albumin assay of cell culture media (**Figure 3E**). There was no remarkable alteration in the gene expression of αSMA between podocytes from the diabetic pmut when compared to diabetic control podocytes, though FSP-1 expression was significantly decreased in diabetic pmut podocytes (**Figure 3F**). Culture media from diabetic pmut podocytes had significantly suppressed levels of IL-1β and IL-6 compared to media from diabetic control podocytes (**Figure 3G**). Podocytes from diabetic pmut mice had no change in lipid uptake; however, lipid oxidation and ATP levels were higher when compared to podocytes from diabetic control mice (**Figure 3H-J**).

Next, podocytes from control mice and pmut mice were co-cultured with isolated control tubules (**Figure 3K**). Interestingly, suppressed levels of TGFβR1/TGFβR2 heterodimerization (DuoLink assay), down-regulated mRNA expression of FSP-1, αSMA and TGFβR1 and mitigated levels of the proinflammatory cytokines IL-1β and IL-6 in tubules cultured with podocytes from diabetic pmut mice were observed (**Figure 3L-N**). The tubules cultured with podocytes from pmut mice did not show any change in lipid uptake, but did show higher levels of lipid oxidation, ATP and miR-29 expression levels (**Figure 3O-Q**). These findings suggest that *Angptl4* deficiency in podocytes protects against EMT in renal tubular cells.

Furthermore, we tested the effect of podocyte *Angptl4* deficiency on the mesenchymal activation of endothelial cells. Podocytes from control mice and pmut mice were co-cultured with control endothelial cells **(Figure S7A)**. Again, we observed down-regulated mRNA expression of FSP-1, αSMA and TGFβR1 and suppressed expression of the proinflammatory cytokines IL-1β and IL-6 in endothelial cells cultured with podocytes from diabetic pmut mice **(Figure S7B-C)**. The endothelial cells cultured with podocytes from pmut mice did not show any change in lipid uptake, but again showed higher levels of lipid oxidation, ATP, and miR-29 expression levels. These cells also showed suppression in permeability as measured by concentration of FITC-dextran **(Figure S7D-G)**. These findings suggest that podocyte *Angptl4* deficiency also protects against EndMT in endothelial cells.

### Angptl4 deficiency in tubules suppresses fibrogenic phenotypes

To further investigate the role of *Angptl4* in tubules, we generated tubule-specific Angtpl4 mutant mice (tmut) by crossing Angptl4^fl/fl^ mice with mice carrying the Tet-inducible *Pax8 Cre* driver (Angptl4^fl/fl^; Tet+; *Pax8 Cre+*) (**Figure 4A**). Tmut mice had significantly suppressed levels of *Angptl4* expression in isolated tubules when compared to control mice **(Figure S7A).** Diabetic tmut mice did not show any remarkable change in body weight or blood glucose levels when compared to diabetic controls (**Figure 4B**). Diabetic tmut mice had decreased kidney weight, lower ACR, decreased area of fibrosis and less glomerulosclerosis when compared to diabetic controls (**Figure 4C**). Ultrastructural analysis by transmission electron microscopy revealed podocyte foot thickening and effacement in diabetic control mice which was less pronounced in diabetic tmut mice (**Figure 4D**).

**Figure 4.**
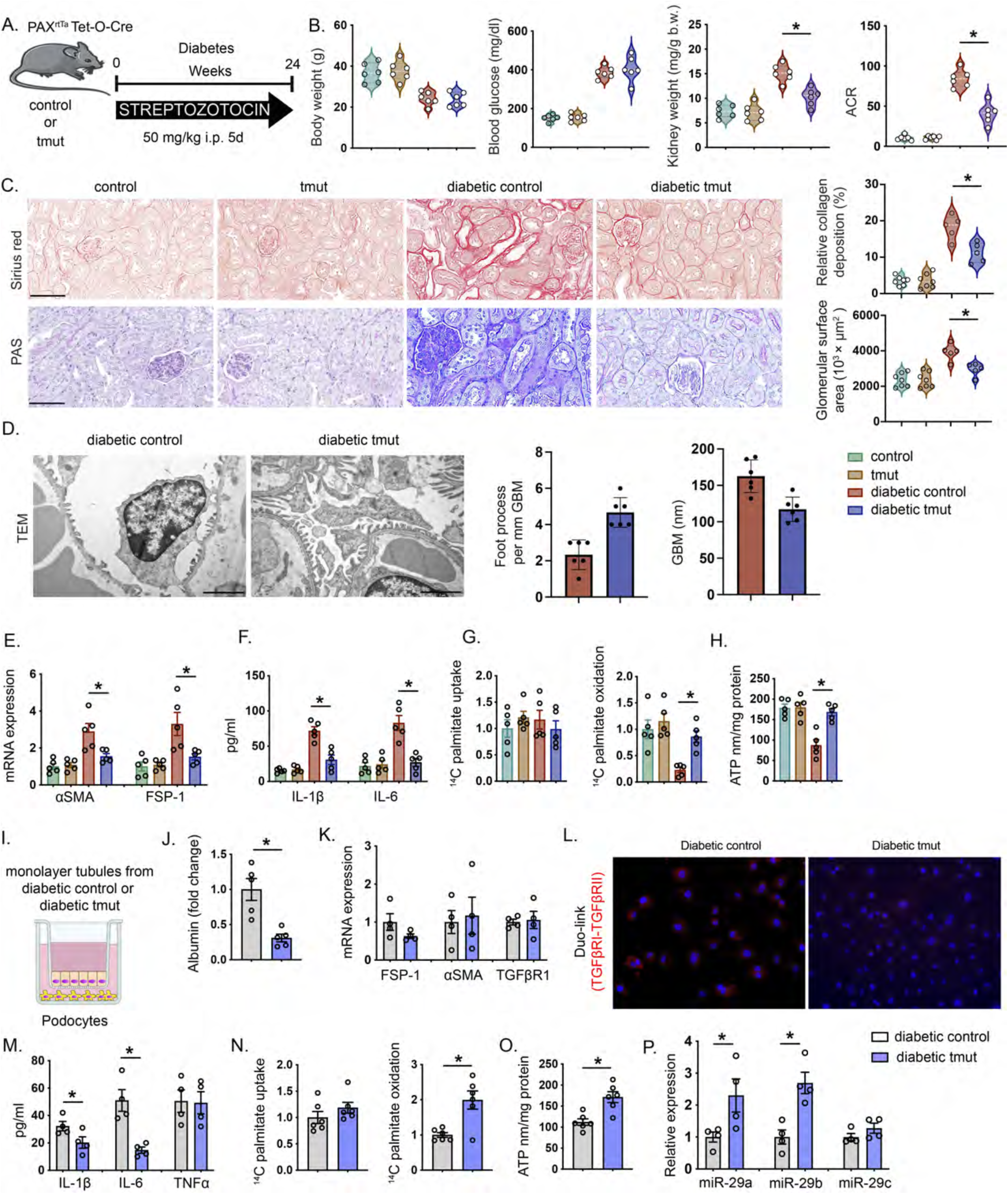
Metabolic reprogramming by tubular Angptl4 loss protects against diabetic kidney disease. **A.** Schematic diagram, showing induction of diabetes in *Angptl4 fl/fl; iPAX8 Cre-* (Control) and in doxycycline-treated *Angptl4 fl/fl; podocin iPax8 Cre+* (tmut). Five doses of STZ (50 mg/kg/day IP) were injected to induce diabetes; after 16 weeks kidneys were evaluated for analysis of fibrosis. Art was created using Servier medical art illustrations. B. Physiological parameters were measured in diabetic and non-diabetic mice of both genotypes. n=5 mice/group combined from two replicate experiments. **C.** Representative histologic images of Sirius red and PAS staining in kidneys from non-diabetic and diabetic control and tmut mice. (Original magnification 300X). Relative area of fibrosis (%) and relative collagen (%) were measured using ImageJ. Scale bar 50 µm. C. Representative transmission electron microscopy images. n=3/group. Scale bar 1 μm. Relative density of podocyte foot processes and glomerular basement membrane (GBM) thickness were calculated using ImageJ software. Six independent images of the staining were analyzed. **E.** qPCR gene expression of α*SMA* and *FSP-1* in tubules from nondiabetic control and tmut and diabetic control and tmut mice. n=5/group. **F.** IL-1β and IL-6 measurement in the media of cultured tubules from nondiabetic control and tmut and diabetic control and tmut mice. n=5/group. **G.** Radiolabeled [C^14^]palmitate uptake analysis and radiolabeled [^14^C]palmitate oxidation were measured in tubules from the indicated groups. CPM of each sample was counted. **H.** ATP measurement in the cultured tubules from nondiabetic control and tmut and diabetic control and tmut mice. **I.** Schematic diagram showing co-culture experimental set-up. Isolated tubules from diabetic control and diabetic tmut mice were cocultured with podocytes from control mice. **J.** After 48 h, albumin was measured in the cell media of podocytes co-cultured either with tubules from diabetic control and tmut mice. **K.** qPCR gene expression analysis of *FSP-1*, α*SMA* and *TGF*β*R1* in the podocytes cocultured either with tubules from diabetic control or diabetic tmut mice **L.** Representative Duolink *In Situ* images of TGF-βR1/2 from podocytes that were co-cultured with tubules either from diabetic control or diabetic tmut mice and corresponding quantification. Combination sites of TGF-βR1/2 in cells were counted in 10 different areas per condition (original magnification 400X). **M.** Measurement of IL-1β, IL-6, and TNFα levels in the cell supernatant. **N.** Radiolabeled [^14^C]palmitate uptake and oxidation, were measured in indicated groups. CPM of each sample was counted. **O.** ATP measurement in the podocytes which were co-cultured with tubules either from diabetic control or diabetic tmut mice. **P.** qPCR gene expression analysis of miR-29a-3p, miR-29b-3p and miR-29c-3p in podocytes which were cocultured with tubules from diabetic control and diabetic tmut mice. Data are mean ± SEM. One-way Anova with Tukey’s multiple comparison test was used to calculate statistical significance. Significance-\**p*<0.05.

Tubules from diabetic tmut mice had suppressed FSP-1 and αSMA gene expression (**Figure 4E**). Culture media from tmut tubules mice had significantly suppressed levels of IL-1β and IL-6 (**Figure 4F**). Isolated tubules from diabetic tmut mice had no change in lipid uptake but did show higher lipid oxidation and ATP levels when compared to tubules from diabetic control mice (**Figure 4G-H**). Tubules from control mice and tmut mice were then co-cultured with control podocytes (**Figure 4I**). Podocytes that were co-cultured with diabetic tmut mice tubules had suppressed albumin permeability as assayed in the culture media (**Figure 4J**) and also did not show any significant change in mRNA expression of FSP-1, αSMA and TGFβR1. However, these podocytes did show suppressed levels of TGFβR1 and TGFβR2 heterodimerization as assessed by DuoLink assay, and lower levels of IL-1β and IL-6 in culture media (**Figure 4K-M**) There was no remarkable change in lipid uptake; however, these podocytes had higher levels of lipid oxidation, ATP and miR-29a and miR-29b expression levels when compared to podocytes cultured with control tubules (**Figure 4N-P**). These findings suggest that *Angptl4* deficiency in tubules protects against podocyte damage in diabetes.

Furthermore, we tested the effect of tubular *Angptl4* deficiency on mesenchymal activation in endothelial cells. Tubules from control mice and tmut mice were co-cultured with control endothelial cells **(Figure S8A)**. We observed lower mRNA expression of FSP-1, αSMA and TGFβR1 as well as suppressed levels of IL-1β and IL-6 in the media from endothelial cells cultured with tubules from diabetic tmut mice compared to those cultured with control tubules **(Figure S8B-C)**. Endothelial cells cultured with tubules from tmut mice did not show any change in lipid uptake, but again showed higher levels of FAO and ATP **(Figure S8D-E)**. Endothelial permeability was suppressed, while miR-29 expression was increased in these cells **(Figure S8F-G)**. Taken together, these findings suggest that tubular *Angptl4* deficiency protects against mesenchymal activation in endothelial cells.

### Podocyte-and tubule-secreted Angptl4 interact with Integrin **β**1 and influence dipeptidyl peptidase-4 (DPP-4)-Integrin **β**1 interactions

Active TGFβ signaling and elevated DPP-4 levels are causative pathways which accelerate renal fibrosis in diabetes^51^. TGFβ1 stimulates DPP-4-Integrin β1 associated fibrogenesis in diabetic endothelium^51^. Duo-link In Situ proximity ligation assay demonstrated that TGFβ1 stimulation increased the interaction of Angptl4 with Integrinβ1 in high glucose-treated HK-2 cells and podocytes (**Figure 5A-B**). To investigate the mechanism underlying the interaction between Angptl4 and Integrinβ1, we analyzed the protein interaction of DPP-4 and Integrin β1, which is critical for mesenchymal activation signal transduction. We observed further proximity between DPP-4 and Integrinβ1 in the tubules which were co-cultured with diabetic pmut mice podocytes and in podocytes which were co-cultured with diabetic tmut tubules when compared to respective diabetic controls, suggesting that either podocyte-or tubule-specific Angptl4 plays a critical role in the DPP-4-Integrinβ1 interaction-mediated mesenchymal signal transduction (**Figure 5C-D**). To test the Angptl4-associated pathogenic roles of DPP-4 and Integrin β1, we treated diabetic control mice with kidney fibrosis with linagliptin, a DPP-4 inhibitor, and neutralizing antibody to Integrinβ1. Both linagliptin and Integrinβ1 neutralization reversed the fibrotic phenotypes and significantly suppressed *Angptl4* expression in diabetic kidneys (**Figure 5 E-F**; **Figure S9A-B)**. To investigate why DPP-4 and Angptl4 levels were elevated in diabetes, we performed miRNA array analysis in control and diabetic mice. miR-29 family members have emerged as key players which are associated with targeting DPP-4 mRNA in endothelial cells in diabetes^51^. Of note, miR-29s have conserved site binding sequences at 3’UTR of Angplt4 mRNA (**Figure 5G**). Luciferase assay analysis revealed that inhibiting miR-29 using an antagomir elevated the Angptl4 3’UTR luciferase activity while use of a mimic or miR-29 overexpression suppressed the luciferase activity of Angptl4 3’UTR (**Figure 5H-I**). microRNA array analysis revealed suppressed expression of miR-29 family members in the kidneys of diabetic mice when compared to those of non-diabetic control mice (**Figure 5J**). Diabetic mice subjected to miR-29 inhibition, through administration of LNA-miR-29, had higher kidney weight and albumin-to-creatinine ratios and significantly worsened fibrosis **(Figure 5K, Figure S10A)**. Suppression of all miR-29s with LNA administration was confirmed by relative gene expression assays **(Figure S10B)**. miR-29 inhibition in UUO-operated mice showed more fibrosis when compared to control-LNA (**Figure 5L**). Treatment with LNA-miR-29 suppressed FAO levels (**Figure 5** **M**) and elevated *Angptl4* gene expression levels (**Figure 5N**). These data suggest that suppression of miR-29 is a key factor which contributes to higher DPP-4 and Angptl4 levels, which in turn interact with integrin β1 and promote mesenchymal activation in tubules and podocytes leading to dysfunction.

**Figure 5.**
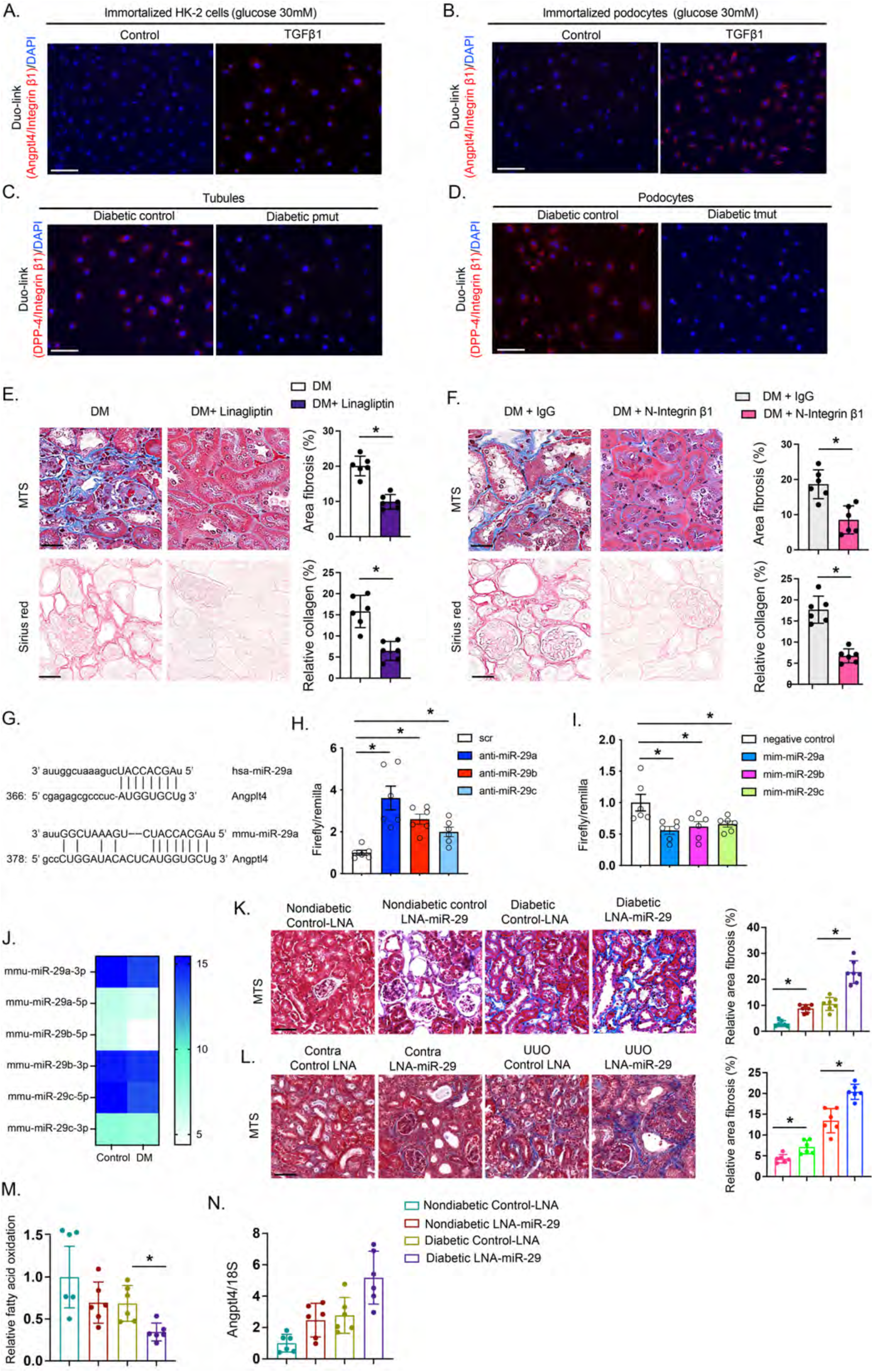
Angplt4 regulates DPP-4/β-integrin interactions. **A.** Proximity between Angptl4 and β-integrin was assessed by DuoLink In Situ Assay in cultured HK-2 cells treated with TGFβ1 (10 ng/ml) for 48 h. For each slide, images at 400X original magnifications were obtained from six different areas. Scale bar 50 µm. **B.** Proximity between Angptl4 and β-integrin was assessed by DuoLink In Situ Assay in immortalized podocytes treated with TGFβ1 (10 ng/ml) for 48 h. For each slide, images at 400X original magnification were obtained from six different areas. Scale bar 50 µm **C.** Proximity between DPP-4 and β-integrin was assessed by DuoLink In Situ Assay in tubules co-cultured either with podocytes from diabetic control or diabetic pmut mice. For each slide, images at 400X original magnification were obtained from six different areas. Scale bar 50 µm. **D.** Proximity between DPP-4 and β-integrin was assessed by DuoLink In Situ Assay in podocytes co-cultured either with tubules from diabetic control or diabetic tmut mice. For each slide, images at 400X original magnification were obtained from six different areas. Scale bar 50 µm. **E.** Representative Masson trichrome and Sirius red staining in kidneys of control and linagliptin-treated diabetic mice. Linagliptin was administered for 8 weeks. Scale bar 50 µm. n=6 mice/group. **F.** Representative Masson trichrome and Sirius red staining in kidneys from IgG control-treated and N-integrin-β1-treated diabetic mice IgG and N-integrin-β1 were provided to diabetic mice for 4 weeks. Scale bar 50 µm. n=6 mice/group. **G.** Data from microRNA.org suggests miR-29a nucleotide sequence alignment with the 3’untranslated region of Angptl4 mRNA. **H.** Angptl4 3’UTR transcriptional activity measurement in the presence of antagomirs for microRNA 29a, b and c. **I.** Angptl4 3’UTR transcriptional activity measurement in the presence of mimics of microRNA 29a, b and c. Two independent experiments were analyzed. **J.** microRNA array analysis of miR-29 family members in control and diabetic mice. **K.** Representative images of Masson trichrome staining of kidneys after LNA-miR-29 treatment in control and diabetic mice. Scale bar 50 µm. n=7 mice/group. **L.** Representative images of Masson trichrome staining in kidneys after LNA-miR-29 treatment in control and unilateral urinary obstruction (UUO)-operated mice. Scale bar 50 µm. n=5 mice/group. **M.** Radiolabeled [^14^C]palmitate oxidation and [^14^C]CO_2_ ptake were measured in indicated groups. CPM of each sample was counted. n=6/group. Data are mean ± SEM. One-way ANOVA was used for the analysis of statistical significance. Significance-\**p*<0.05.

Clement et al., found a pro-proteinuric form of Angptl4 in glomeruli which is hyposialylated^37^. Conversion of Angplt4 from the hyposialylated to the sialylated form by treatment with N-acetyl-D-mannosamine suppressed the level of proteinuria and disease phenotype in this mouse model of nephrotic syndrome. Therefore, we tested the sialic acid precursor N-acetyl mannosamine (NAM) in our mouse model of diabetic kidney disease. NAM did not cause any differences in body weight, blood glucose or blood pressure but significantly suppressed albuminuria and kidney weight. NAM treatment significantly suppressed fibrosis, collagen deposition and glomerulosclerosis in diabetic mice **(Figure S11)**.

### Kidney-specific Angptl4 ASO improves the fibrogenic phenotype in diabetic mice

We also analyzed the effects of some well-known and effective drugs, such as dichloroacetate (DCA), which causes increased conversion of pyruvate to acetyl-CoA, the glycolysis inhibitor 2-deoxy-glucose (2-DG), the angiotensin-converting enzyme inhibitor (ACEi) imidapril, and the lipid lowering drugs fenofibrate and simvastatin, on *Angptl4* expression in diabetic kidneys. The anti-inflammatory peptide N-acetyl-seryl-aspartyl-lysyl-proline (Ac-SDKP) was also used, both alone and in combination with ACEi. DCA, 2-DG, Ac-SDKP, Ac-SDKP +ACEi, ACEi and fenofibrate significantly suppressed fibrosis and *Angptl4* expression suggesting that blockade of kidney-specific *Angptl4* could be an important strategy in the management of diabetic kidney disease **(Figure S12)**.

Data from human genetic studies have demonstrated that loss-of-function mutations in the *Angptl4* locus are linked with reduced type 2 diabetes and risk of cardiovascular disease^48,60–64^. Given the improved kidney phenotype observed in mice lacking *Angptl4* expression in podocytes and tubules, we evaluated targeted inhibition of *Angptl4* expression. We treated diabetic mice with a kidney-specific anti-sense oligonucleotide (ASO) against Angptl4 which has high affinity for the kidney cortex. Diabetic mice were injected subcutaneously with Angptl4 ASO or control ASO for eight weeks (**Figure 6A**). After treatment, *Angptl4* mRNA was significantly suppressed in both nondiabetic and diabetic mice (**Figure 6B**). Angptl4 ASO did not cause any change in body weight, blood glucose or blood pressure in either group. However, Angptl4 ASO caused significant reduction in the kidney weight and ACR of diabetic mice (**Figure 6C-G**).

**Figure 6.**
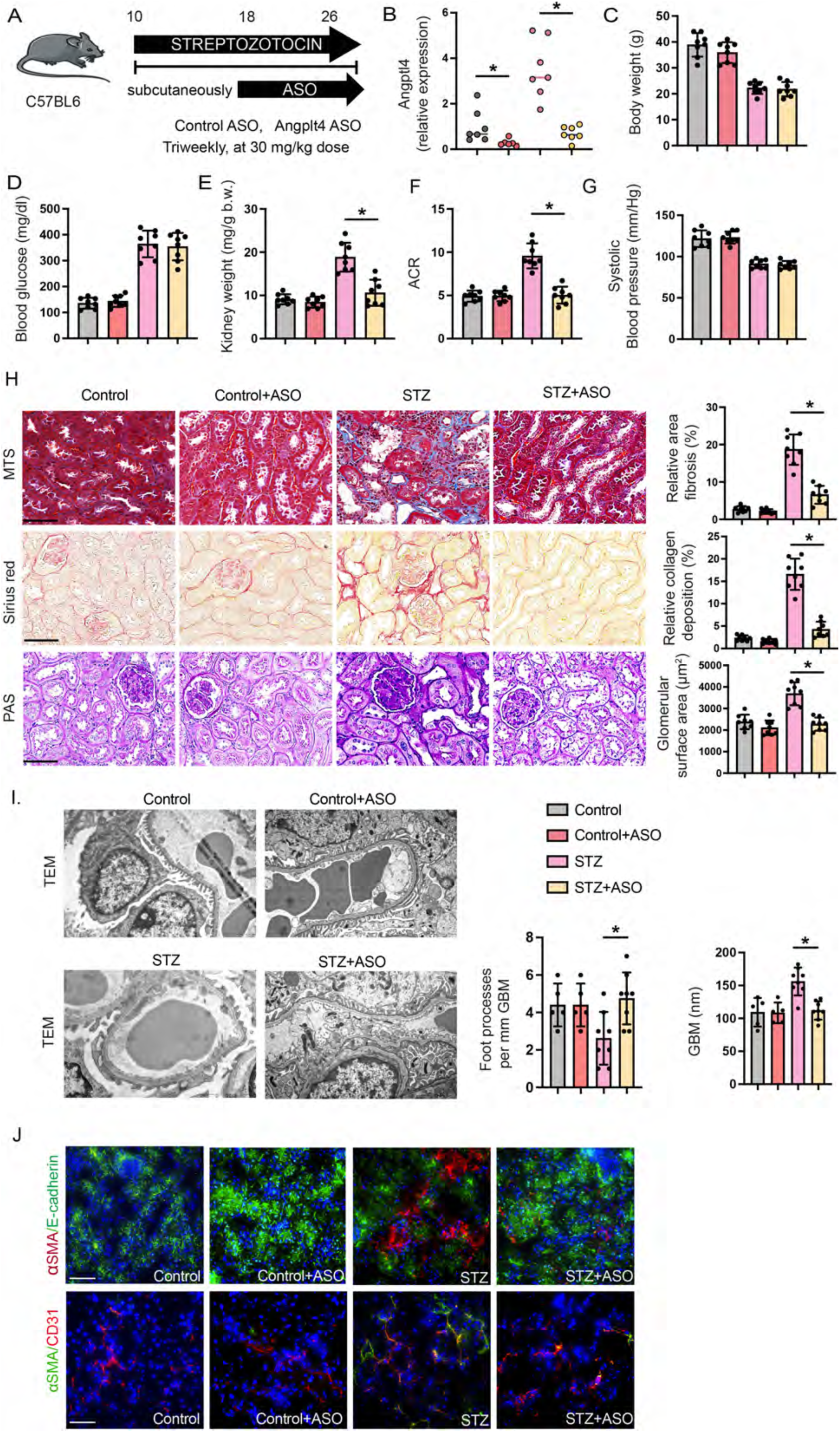
Kidney-specific ASO treatment protects against diabetic kidney disease. **A.** Schematic diagram showing the treatment protocol of control ASO and kidney-specific Angplt4 ASO treatment in diabetic and non-diabetic C57BL/6 mice. Five doses of STZ (50 mg/kg/day IP) were injected to induce diabetes; after 16 weeks the mice received either control ASO or Angptl4 ASO for eight weeks. Art was created using Servier medical art illustrations. **B.** RT-qPCR gene expression analysis of *Angptl4* in kidneys of nondiabetic and diabetic, control ASO-and Angplt4 ASO-treated mice. n=7/group. **C-G.** Physiological parameters of nondiabetic and diabetic, control ASO-and Angplt4 ASO-treated mice. n=8 mice/group, combined from two replicate experiments. **H.** Representative histologic images from MTS, Sirius red and PAS staining in kidneys from nondiabetic and diabetic, control ASO-and Angplt4 ASO-treated mice (original magnification 300X). Relative area of fibrosis (%) and relative collagen (%) were quantified using ImageJ. Scale bar 50 µm. **I.** Representative transmission electron microscopy images are shown. n=3/group. Scale bar 1 μm. Relative density of podocyte foot processes and glomerular basement membrane (GBM) thickness were calculated using ImageJ software. Six independent images of the staining were analyzed. **J.** Immunofluorescence analysis of αSMA/E-cadherin and αSMA/CD31 in kidneys of ASO treatment in control and diabetic mice. FITC-labeled E-cadherin, rhodamine-labeled αSMA and DAPI (nuclei, blue); FITC-labeled αSMA, rhodamine-labeled CD31 and DAPI (nuclei, blue), were used. Representative merged (original magnification 400X) images are shown. Scale bar 50 μm. n=8 mice/group. Three independent experiments were analyzed. Data are mean ± SEM. One-way ANOVA was used for the analysis of statistical significance. Significance-\**p*<0.05.

Angptl4 ASO did not cause remarkable alterations in non-diabetic control mice, but caused significant reduction in fibrosis, collagen deposition and glomerulosclerosis in diabetic mice, suggesting that Angptl4 ASO improves the kidney phenotype (**Figure 6H**). Electron microscopy revealed partial restoration of normal podocyte structure and GBM caliber in ASO-treated diabetic mice when compared to vehicle-treated diabetic controls (**Figure 6I**). Diabetes enhanced the rate of EndMT and EMT in the kidneys when compared to nondiabetic mice as evidenced by higher co-localization of αSMA in E-cadherin-positive and CD31-positive cells. This effect was significantly suppressed in the Angptl4 ASO-treated kidneys, which approximated the ASO-treated non-diabetic kidneys (**Figure 6J**).

## Discussion

This study shows the crucial role of podocyte and tubular *Angptl4* in the fibrogenic process of diabetic kidney disease. Our results demonstrate that these cell-type specific *Angptl4* are key fibrogenic molecules, activating DPP-4-Integrin β1 signaling and its linked mesenchymal transdifferentiation processes in diabetic kidneys. Further, these results demonstrate how two components of this disease, fibrosis and mesenchymal metabolic shifts, are linked in the tubules and podocytes. *Angptl4* is expressed in kidneys, however its roles there have been incompletely investigated^65^. Human studies have shown a positive correlation between Angptl4 levels and expression of characteristics of diabetic nephropathy^66^. In cultured glomerular mesangial cells, Angptl4 deficiency inhibits both the inflammatory response and extracellular matrix accumulation^67^.

*Angptl4* is a known inhibitor LPL, an enzyme that catalyzes the hydrolysis of triglycerides into fatty acids, which are utilized by peripheral tissues and kidney tubules^65^. Our data demonstrate that *Angptl4* expression is upregulated, and LPL expression and its associated activities are downregulated, in fibrotic kidneys during diabetes. However, while chemical inhibition of LPL in diabetic mice caused an elevation in plasma triglycerides, it did not alter renal fibrosis, suggesting that LPL is not a key molecule in the development of diabetic kidney fibrosis. Analysis of diabetic kidneys from Angptl4 mutant mice shows suppression of renal fibrosis, proteinuria, pro-inflammatory cytokines, and mesenchymal activation in tubules and endothelial cells when compared to diabetic controls. These results suggest that *Angptl4* is associated with fibrosis, proteinuria, inflammation and mesenchymal activation in diabetic kidneys. It is clear from these data that Angptl4 is one of the catalysts of renal fibrosis in diabetes and leads to disruption of cytokine and chemokine homeostasis by up regulating TGFβ signaling. These processes alter metabolic homeostasis by predisposing towards defective fatty acid metabolism and associated mesenchymal activation in tubules and endothelial cells.

These observations implicate renal Angptl4 as a mediator and potential therapeutic target for the treatment of diabetic kidney disease. However, a clear role for Angplt4 in regulating whole body lipid and glucose metabolism has not been fully elucidated due to lack of availability of relevant tissue-specific knock out mouse models. Severe systemic metabolic complications such as gut inflammation and ascites have limited investigators’ ability to analyze the beneficial functions of Angptl4 deficiency in diabetes^68^. To capitalize on the specificity of Angptl4 expression in the kidney, we generated podocyte-specific Angplt4 mutant and tubule-specific mutant mice. Clement and colleagues demonstrated the critical role of podocyte-specific Angptl4 in causing proteinuria in nephrotic syndrome^37^. Angplt4 was found to express in two forms, the sialylated (normal) form and the hyposialylated (abnormal) form. The hyposialylated form is causative for proteinuria. Conversion of the hyposialylated to the sialylated form by treatment with N-acetyl-D-mannosamine was noted to significantly suppress the levels of proteinuria in a mouse model of nephrotic syndrome^37^. Clement et al., also demonstrated that circulatory sialylated Angplt4 suppresses proteinuria while increasing hypertriglyceridemia in mice^38^. In line with previous findings, our data demonstrate that N-acetyl-D-mannosamine treatment in diabetic mice suppressed fibrosis and proteinuria and restored kidney structure in diabetic mice. Further, our data show that blocking Angptl4 in podocytes and tubules effectively suppresses proteinuria, glomerular fibrosis and interstitial fibrosis in diabetic mice. One limitation of our study is that we could not demonstrate whether Angptl4 in diabetic kidneys exists in the hyposialylated or sialylated form due to the lack of a suitable mouse Angplt4 antibody, thus precluding us from evaluating Angptl4 at the protein level. However, based on previous and current data, we suspect that diabetes may cause renal secretion of the hyposialylated form of Angptl4 protein.

To investigate how Angptl4 accelerates mesenchymal activation, we analyzed FAO in tubular epithelial cells, endothelial cells, and podocytes. Defective FAO in tubular epithelial cells is a key process in the development of myofibroblasts and fibrogenesis in diabetic kidneys^41^. Kidney tubules are enriched in mitochondria due to their high ATP demand and therefore are highly dependent on FAO for ATP, since oxidation of fatty acid generates far more ATP than oxidation of glucose. Our data suggest that upregulation of *Angptl4* expression is associated with increased fibrosis and suppression of FAO. *Angptl4* deficiency resulted in increased FAO and decreased abnormal glucose metabolism. Metabolic reprogramming and cytokine reprogramming by *Angplt4* deficiency is a crucial step in the anti-mesenchymal and anti-fibrotic mechanisms occurring in kidney tubules during diabetes.

To begin to understand how Angptl4 regulates FAO and defective glycolysis, we analyzed DPP-4-Integrin β1 signaling levels and the associated TGFβ dimerization which is the first step in activating the TGFβ signaling cascade. DPP-4 is a well-known molecule influencing TGFβ-associated EndMT and fibrosis in endothelial cells^51,55,69^. Proximity ligation assays revealed that TGFβ1-associated fibrogenesis is linked with closer Integrin β1 and Angptl4 proximity suggesting that Angptl4 interacts with Integrin β1 and influences activation of the mesenchymal signal cascade in tubules. Podocyte-secreted Angptl4 and tubule-secreted Angptl4 influence the DPP-4-Integrin β1 interaction and TGFβR1 and TGFβR2 dimerization, resulting in a cascade of signaling processes mediated through Smad proteins, primarily Smad3. Phosphorylated Smad3 alters the cytoplasmic pathway and also enters the nucleus where it represses the transcription of genes responsible for FAO and induces the transcription of genes related to fibrogenesis in tubules^41^. In podocytes, higher levels of the DPP-4-Integrin β1 signaling cascade cause loss of glomerular basement membrane, alterations in podocyte cell permeability, changes in foot process effacement and podocyte cell death. These data indicate that podocyte-and tubule-Angptl4 interact with Integrin β1 and promote DPP-4-Integrin β1 interaction and TGFβ signaling, and that these cumulative effects lead to associated activation of mesenchymal signal transduction in tubules and endothelial cells, foot process effacement, increased albuminuria and podocyte death.

To further investigate the interaction between DPP-4-Integrin β1 and Angptl4, we analyzed microRNA array data from control and TGFβ-treated cultured tubular epithelial cells. Data suggest miR-29-3p is critical in regulating *Angptl4* expression levels and targets the 3’UTR of its mRNA. miR-29-3p targets the 3’UTR of DPP-4 mRNA in endothelial cells and regulates EndMT and fibrogenesis^51^. From our data, miR-29b-3p deficiency in TGFβ-treated high-glucose stimulated tubules and in diabetic tubules is identified as a key regulator of DPP-4 and Angptl4 levels. miR-29b-3p targets DPP-4 and Angptl4 levels in the tubules. LNA-miR-29 treatment has been shown to promote favorable plaque remodeling in atherosclerotic mice^70^, worsen the features of fibrosis, derange kidney structure and cause severe proteinuria in mouse models of diabetic kidney disease; however, miR-29b was found to be reno-protective in *db/db* mice^71^. These observations suggest that miR-29-3p is a crucial regulator of DPP-4 and Angptl4 and influence fibrogenesis in diabetic kidneys.

Past studies have shown Angptl4 to be a potential metabolic regulator which is linked with several metabolic diseases^72–74^. In one study, Dewey et al., used a neutralizing antibody to Angptl4 in both humanized mice and non-human primates which resulted in severe side effects and systemic metabolic abnormalities^60^. Our data using whole body Angptl4 mutant mice, podocyte-specific and tubule-specific mutant mice suggest that Angptl4 has promising drug targetability against diabetic kidney disease. Hence, we evaluated a kidney-specific ASO in our model of diabetic kidney disease. This design is advantageous in that it avoids the unfavorable systemic effects of whole body Angptl4 loss. The kidney specific ASO was designed to have the highest affinity for the cortex and limited affinity for the medulla. Our results demonstrate that Angptl4 ASO-treated mice have partially restored kidney structure, substantially suppressed glomerular and cortical fibrosis and improved proteinuria, compared to control ASO-treated mice, without affecting body weight, blood glucose or blood pressure.

Podocyte-and tubule-Angpl4 are key fibrogenic molecules and their loss is protective against diabetic kidney disease and fibrogenesis by metabolic reprogramming which is driven by suppression in DPP-4-Integrin β1 signaling and TGFβ signaling. Kidney-specific Angplt4 ASO reversed the fibrogenic phenotypes of diabetic kidney disease. Taken together, these data add significant information to the understanding of Angplt4 biology and offer an exciting possibility as a new therapeutic option for the treatment of this condition.

## Acknowledgements

This work is supported by the following grants from the National Institutes of Health: R01HL144476 (AD), R01HL162580 (AD). R01DK133143 (GIS), RC2DK120534 (GIS), UC2DK134901 (GIS), P30DK045735 (GIS), R35HL135820 (CFH) and R01HL131952 (JEG). KK is supported by a grant from the Japan Society for the Promotion of Science (22K08330) and a grant from The Ministry of Health Labour and Welfare (202112004A). KK is under a consultancy agreement with Boehringer Ingelheim. KK collaborated with Boehringer Ingelheim, Taisho Pharma and Kowa for a project not related to this manuscript. KK’s department at Shimane University was supported by funds from Boehringer Ingelheim, Mitsubishi Tanabe Pharma, Taisho Pharmaceutical, Ono Pharmaceutical, Bayer, Kowa, Nipro, and Life Scan Japan. KK received lecture honoraria from Dainippon-Sumitomo Pharma, Astellas, Astra Zeneca, Ono, Otsuka, Taisho, Tanabe-Mitsubishi, Eli Lilly, Boehringer-Ingelheim, Novo Nordisk, Bayer, Sanofi, and Kowa. KK is the recipient of Japan Diabetes Society Carrier Development Award supported by Sanofi. We thank William Sessa for use of the LNA-miR-29.

## Author’s contribution

SPS led the project, proposed the original idea and experimental design and wrote the paper. SPS, HZ, RS, MM, LG, OS and BA performed the experiments. BKR performed the bioinformatics analysis. KK, DK, AD, CFH and GIS provided intellectual guidance. CFH and WS also provided reagents and animals. DK and KK guided some experiments and provided intellectual output. JEG supervised the project, validated the data, made intellectual contributions, and performed final editing the manuscript.

## References

1 Thomas, M. C., et al. Diabetic kidney disease. Nat Rev Dis Primers 1, 15018 (2015). 10.1038/nrdp.2015.18

2 Gnudi, L., Coward, R. J. M. & Long, D. A. Diabetic Nephropathy: Perspective on Novel Molecular Mechanisms. Trends Endocrinol Metab 27, 820–830 (2016). 10.1016/j.tem.2016.07.002

3 Held, P. J. et al. The United States Renal Data System’s 1991 annual data report: an introduction. Am J Kidney Dis 18, 1–16 (1991).

4 Cooper, M. & Warren, A. M. A promising outlook for diabetic kidney disease. Nat Rev Nephrol 15, 68–70 (2019). 10.1038/s41581-018-0092-5

5. Srivastava, S. P. & Kanasaki, K. Editorial: Receptor Biology and Cell Signaling in Diabetes. Front Pharmacol 13, 864117 (2022). 10.3389/fphar.2022.864117

6 Srivastava, S. P., Hedayat, F. A., Kanasaki, K. & Goodwin, J. E. microRNA Crosstalk Influences Epithelial-to-Mesenchymal, Endothelial-to-Mesenchymal, and Macrophage-to-Mesenchymal Transitions in the Kidney. Front Pharmacol 10, 904 (2019). 10.3389/fphar.2019.00904

7 Falke, L. L., Gholizadeh, S., Goldschmeding, R., Kok, R. J. & Nguyen, T. Q. Diverse origins of the myofibroblast-implications for kidney fibrosis. Nat Rev Nephrol 11, 233–244 (2015). 10.1038/nrneph.2014.246

8 Meng, X. M., Nikolic-Paterson, D. J. & Lan, H. Y. Inflammatory processes in renal fibrosis. Nat Rev Nephrol 10, 493–503 (2014). 10.1038/nrneph.2014.114

9 Lovisa, S. et al. Epithelial-to-mesenchymal transition induces cell cycle arrest and parenchymal damage in renal fibrosis. Nat Med 21, 998–1009 (2015). 10.1038/nm.3902

10 Zeisberg, M. & Neilson, E. G. Mechanisms of tubulointerstitial fibrosis. Journal of the American Society of Nephrology : JASN 21, 1819–1834 (2010). 10.1681/ASN.2010080793

11 Srivastava, S. P. et al. Endothelial SIRT3 regulates myofibroblast metabolic shifts in diabetic kidneys. iScience 24, 102390 (2021). 10.1016/j.isci.2021.102390

12 Allison, S. J. Fibrosis: Targeting EMT to reverse renal fibrosis. Nat Rev Nephrol 11, 565 (2015). 10.1038/nrneph.2015.133

13 Grande, M. T. et al. Snail1-induced partial epithelial-to-mesenchymal transition drives renal fibrosis in mice and can be targeted to reverse established disease. Nat Med 21, 989–997 (2015). 10.1038/nm.3901

14 Li, J. et al. Endothelial FGFR1 (Fibroblast Growth Factor Receptor 1) Deficiency Contributes Differential Fibrogenic Effects in Kidney and Heart of Diabetic Mice. Hypertension 76, 1935–1944 (2020). 10.1161/HYPERTENSIONAHA.120.15587

15 Kuppe, C. et al. Decoding myofibroblast origins in human kidney fibrosis. Nature 589, 281–286 (2021). 10.1038/s41586-020-2941-1

16 Humphreys, B. D. et al. Fate tracing reveals the pericyte and not epithelial origin of myofibroblasts in kidney fibrosis. The American journal of pathology 176, 85–97 (2010). 10.2353/ajpath.2010.090517

17 Yang, J. et al. Guidelines and definitions for research on epithelial-mesenchymal transition. Nat Rev Mol Cell Biol 21, 341–352 (2020). 10.1038/s41580-020-0237-9

18 LeBleu, V. S. & Neilson, E. G. Origin and functional heterogeneity of fibroblasts. FASEB J 34, 3519–3536 (2020). 10.1096/fj.201903188R

19 Lovisa, S., Zeisberg, M. & Kalluri, R. Partial Epithelial-to-Mesenchymal Transition and Other New Mechanisms of Kidney Fibrosis. Trends Endocrinol Metab 27, 681–695 (2016). 10.1016/j.tem.2016.06.004

20 Lovisa, S. Epithelial-to-Mesenchymal Transition in Fibrosis: Concepts and Targeting Strategies. Front Pharmacol 12, 737570 (2021). 10.3389/fphar.2021.737570

21 Meng, X. M., Tang, P. M., Li, J. & Lan, H. Y. TGF-beta/Smad signaling in renal fibrosis. Front Physiol 6, 82 (2015). 10.3389/fphys.2015.00082

22 Meng, X. M., Nikolic-Paterson, D. J. & Lan, H. Y. TGF-beta: the master regulator of fibrosis. Nat Rev Nephrol 12, 325–338 (2016). 10.1038/nrneph.2016.48

23 Huang, S. & Susztak, K. Epithelial Plasticity versus EMT in Kidney Fibrosis. Trends Mol Med 22, 4–6 (2016). 10.1016/j.molmed.2015.11.009

24 Bielesz, B. et al. Epithelial Notch signaling regulates interstitial fibrosis development in the kidneys of mice and humans. J Clin Invest 120, 4040–4054 (2010). 10.1172/JCI43025

25 He, W. et al. Wnt/beta-catenin signaling promotes renal interstitial fibrosis. Journal of the American Society of Nephrology : JASN 20, 765–776 (2009). 10.1681/ASN.2008060566

26 Li, X. et al. Wnt5a promotes renal tubular inflammation in diabetic nephropathy by binding to CD146 through noncanonical Wnt signaling. Cell Death Dis 12, 92 (2021). 10.1038/s41419-020-03377-x

27 Srivastava, S. P. et al. Loss of endothelial glucocorticoid receptor accelerates diabetic nephropathy. Nat Commun 12, 2368 (2021). 10.1038/s41467-021-22617-y

28 Srivastava, S. P. et al. Podocyte Glucocorticoid Receptors Are Essential for Glomerular Endothelial Cell Homeostasis in Diabetes Mellitus. J Am Heart Assoc 10, e019437 (2021). 10.1161/JAHA.120.019437

29 Fabian, S. L. et al. Hedgehog-Gli pathway activation during kidney fibrosis. Am J Pathol 180, 1441–1453 (2012). 10.1016/j.ajpath.2011.12.039

30 Ding, H. et al. Sonic hedgehog signaling mediates epithelial-mesenchymal communication and promotes renal fibrosis. J Am Soc Nephrol 23, 801–813 (2012). 10.1681/ASN.2011060614

31 Rauhauser, A. A. et al. Hedgehog signaling indirectly affects tubular cell survival after obstructive kidney injury. Am J Physiol Renal Physiol 309, F770–778 (2015). 10.1152/ajprenal.00232.2015

32 Edeling, M., Ragi, G., Huang, S., Pavenstadt, H. & Susztak, K. Developmental signalling pathways in renal fibrosis: the roles of Notch, Wnt and Hedgehog. Nat Rev Nephrol 12, 426–439 (2016). 10.1038/nrneph.2016.54

33 Zoja, C., Xinaris, C. & Macconi, D. Diabetic Nephropathy: Novel Molecular Mechanisms and Therapeutic Targets. Front Pharmacol 11, 586892 (2020). 10.3389/fphar.2020.586892

34 Kersten, S. Role and mechanism of action of angiopoietin-like protein ANGPTL4 in plasma lipid metabolism. J Lipid Res, 100150 (2021). 10.1016/j.jlr.2021.100150

35 Morris, A. Obesity: ANGPTL4 - the link binding obesity and glucose intolerance. Nat Rev Endocrinol 14, 251 (2018). 10.1038/nrendo.2018.35

36 Dijk, W. & Kersten, S. Regulation of lipoprotein lipase by Angptl4. Trends Endocrinol Metab 25, 146–155 (2014). 10.1016/j.tem.2013.12.005

37 Clement, L. C. et al. Podocyte-secreted angiopoietin-like-4 mediates proteinuria in glucocorticoid-sensitive nephrotic syndrome. Nat Med 17, 117–122 (2011). 10.1038/nm.2261

38 Clement, L. C. et al. Circulating angiopoietin-like 4 links proteinuria with hypertriglyceridemia in nephrotic syndrome. Nat Med 20, 37–46 (2014). 10.1038/nm.3396

39 Chugh, S. S., Mace, C., Clement, L. C., Del Nogal Avila, M. & Marshall, C. B. Angiopoietin-like 4 based therapeutics for proteinuria and kidney disease. Front Pharmacol 5, 23 (2014). 10.3389/fphar.2014.00023

40 Mizunuma, Y. et al. CD-1(db/db) mice: A novel type 2 diabetic mouse model with progressive kidney fibrosis. J Diabetes Investig 11, 1470–1481 (2020). 10.1111/jdi.13311

41 Kang, H. M. et al. Defective fatty acid oxidation in renal tubular epithelial cells has a key role in kidney fibrosis development. Nat Med 21, 37–46 (2015). 10.1038/nm.3762

42 Srivastava, S. P. et al. SIRT3 deficiency leads to induction of abnormal glycolysis in diabetic kidney with fibrosis. Cell Death Dis 9, 997 (2018). 10.1038/s41419-018-1057-0

43 Zhou, H. L. et al. Metabolic reprogramming by the S-nitroso-CoA reductase system protects against kidney injury. Nature 565, 96–100 (2019). 10.1038/s41586-018-0749-z

44 Schoors, S. et al. Fatty acid carbon is essential for dNTP synthesis in endothelial cells. Nature 520, 192–197 (2015). 10.1038/nature14362

45 Tran, M. T. et al. PGC1alpha drives NAD biosynthesis linking oxidative metabolism to renal protection. Nature 531, 528–532 (2016). 10.1038/nature17184

46 Qi, W. et al. Pyruvate kinase M2 activation may protect against the progression of diabetic glomerular pathology and mitochondrial dysfunction. Nat Med 23, 753–762 (2017). 10.1038/nm.4328

47 Xiong, J. et al. A Metabolic Basis for Endothelial-to-Mesenchymal Transition. Mol Cell 69, 689–698 e687 (2018). 10.1016/j.molcel.2018.01.010

48 Gusarova, V. et al. Genetic inactivation of ANGPTL4 improves glucose homeostasis and is associated with reduced risk of diabetes. Nat Commun 9, 2252 (2018). 10.1038/s41467-018-04611-z

49 Singh, A. K. et al. Hepatocyte-specific suppression of ANGPTL4 improves obesity-associated diabetes and mitigates atherosclerosis in mice. J Clin Invest (2021). 10.1172/JCI140989

50 Srivastava, S. P., Goodwin, J. E., Kanasaki, K. & Koya, D. Metabolic reprogramming by N-acetyl-seryl-aspartyl-lysyl-proline protects against diabetic kidney disease. Br J Pharmacol 177, 3691–3711 (2020). 10.1111/bph.15087

51 Kanasaki, K. et al. Linagliptin-mediated DPP-4 inhibition ameliorates kidney fibrosis in streptozotocin-induced diabetic mice by inhibiting endothelial-to-mesenchymal transition in a therapeutic regimen. Diabetes 63, 2120–2131 (2014). 10.2337/db13-1029

52 Seth, P. P. et al. Design, synthesis and evaluation of constrained methoxyethyl (cMOE) and constrained ethyl (cEt) nucleoside analogs. Nucleic Acids Symp Ser (Oxf), 553–554 (2008). 10.1093/nass/nrn280

53 Srivastava, S. P. & Goodwin, J. E. Loss of endothelial glucocorticoid receptor accelerates organ fibrosis in db/db mice. Am J Physiol Renal Physiol (2023). 10.1152/ajprenal.00105.2023

54 Li, J. et al. FGFR1 is critical for the anti-endothelial mesenchymal transition effect of N-acetyl-seryl-aspartyl-lysyl-proline via induction of the MAP4K4 pathway. Cell Death Dis 8, e2965 (2017). 10.1038/cddis.2017.353

55 Shi, S. et al. Interactions of DPP-4 and integrin beta1 influences endothelial-to-mesenchymal transition. Kidney Int 88, 479–489 (2015). 10.1038/ki.2015.103

56 Kaur, K., Pandey, A. K., Srivastava, S., Srivastava, A. K. & Datta, M. Comprehensive miRNome and in silico analyses identify the Wnt signaling pathway to be altered in the diabetic liver. Mol Biosyst 7, 3234–3244 (2011). 10.1039/c1mb05041a

57 Tian, Z. & Liang, M. Renal metabolism and hypertension. Nat Commun 12, 963 (2021). 10.1038/s41467-021-21301-5

58 Huang, H. & Parikh, S. M. Skimming the fat in diabetic kidney disease: KIM-1 and tubular fatty acid uptake. Kidney Int 100, 969–972 (2021). 10.1016/j.kint.2021.06.038

59 Ralto, K. M., Rhee, E. P. & Parikh, S. M. NAD(+) homeostasis in renal health and disease. Nat Rev Nephrol 16, 99–111 (2020). 10.1038/s41581-019-0216-6

60 Dewey, F. E. et al. Inactivating Variants in ANGPTL4 and Risk of Coronary Artery Disease. N Engl J Med 374, 1123–1133 (2016). 10.1056/NEJMoa1510926

61 Klarin, D. et al. Genetics of blood lipids among ∼300,000 multi-ethnic participants of the Million Veteran Program. Nat Genet 50, 1514–1523 (2018). 10.1038/s41588-018-0222-9

62 Mooradian, A. D. Dyslipidemia in type 2 diabetes mellitus. Nat Clin Pract Endocrinol Metab 5, 150–159 (2009). 10.1038/ncpendmet1066

63 Do, R. et al. Common variants associated with plasma triglycerides and risk for coronary artery disease. Nat Genet 45, 1345–1352 (2013). 10.1038/ng.2795

64 Barja-Fernandez, S. et al. Plasma ANGPTL-4 is Associated with Obesity and Glucose Tolerance: Cross-Sectional and Longitudinal Findings. Mol Nutr Food Res 62, e1800060 (2018). 10.1002/mnfr.201800060

65 Scerbo, D. et al. Kidney triglyceride accumulation in the fasted mouse is dependent upon serum free fatty acids. J Lipid Res 58, 1132–1142 (2017). 10.1194/jlr.M074427

66 Al Shawaf, E. et al. ANGPTL4: A Predictive Marker for Diabetic Nephropathy. J Diabetes Res 2019, 4943191 (2019). 10.1155/2019/4943191

67 Qin, L., Zhang, R., Yang, S., Chen, F. & Shi, J. Knockdown of ANGPTL-4 inhibits inflammatory response and extracellular matrix accumulation in glomerular mesangial cells cultured under high glucose condition. Artif Cells Nanomed Biotechnol 47, 3368–3373 (2019). 10.1080/21691401.2019.1649274

68 Lichtenstein, L. et al. Angptl4 protects against severe proinflammatory effects of saturated fat by inhibiting fatty acid uptake into mesenteric lymph node macrophages. Cell Metab 12, 580–592 (2010). 10.1016/j.cmet.2010.11.002

69 Scheen, A. J. Pharmacokinetics of dipeptidylpeptidase-4 inhibitors. Diabetes Obes Metab 12, 648–658 (2010). 10.1111/j.1463-1326.2010.01212.x

70 Ulrich, V. et al. Chronic miR-29 antagonism promotes favorable plaque remodeling in atherosclerotic mice. EMBO Mol Med 8, 643–653 (2016). 10.15252/emmm.201506031

71 Chen, H. Y. et al. MicroRNA-29b inhibits diabetic nephropathy in db/db mice. Mol Ther 22, 842–853 (2014). 10.1038/mt.2013.235

72 Dai, L. et al. Weighted Gene Co-Expression Network Analysis Identifies ANGPTL4 as a Key Regulator in Diabetic Cardiomyopathy via FAK/SIRT3/ROS Pathway in Cardiomyocyte. Front Endocrinol (Lausanne) 12, 705154 (2021). 10.3389/fendo.2021.705154

73 Spitler, K. M., Shetty, S. K., Cushing, E. M., Sylvers-Davie, K. L. & Davies, B. S. J. Chronic high-fat feeding and prolonged fasting in liver-specific ANGPTL4 knockout mice. Am J Physiol Endocrinol Metab 321, E464–E478 (2021). 10.1152/ajpendo.00144.2021

74 Spitler, K. M., Shetty, S. K., Cushing, E. M., Sylvers-Davie, K. L. & Davies, B. S. J. Regulation of plasma triglyceride partitioning by adipose-derived ANGPTL4 in mice. Sci Rep 11, 7873 (2021). 10.1038/s41598-021-87020-5

