## Supplementary Figure Legends for "Renal Angptl4 is a key fibrogenic molecule in progressive diabetic kidney disease"

**Figure S1**

mRNA array analysis in control and diabetic mice. Gene expression networking was analyzed using STRING (<https://string-db.org/>).

**Figure S2**

KEGG Pathway analysis, Gene Ontology (GO)-molecular function and GO- biological process analysis from the mRNA array data set.

**Figure S3**

**A.** Sirius red and PAS staining in kidneys of poloxamer-injected and vehicle-injected diabetic mice. Representative histologic images are shown (original magnification 300X). Relative area of fibrosis (%) was measured using ImageJ. Scale bar 50 µm. n=5/group. **B.** Plasma triglyceride measurement through calorimetry. n=5/group. **C.** Lipoprotein lipase (LPL) measurement. n=5/group. Data are mean ± SEM. Student t-test (unpaired two-tailed) was used for analysis of statistical significance. Significance- *p<0.05.

**Figure S4**

A. Relative gene expression analysis of Angplt3 and Angptl4 by qPCR in low-glucose (5.55 mM)- and high-glucose- (30 mM) treated cultured HK-2 cells. Data are mean ± SEM. Student t-test (unpaired two-tailed) was used for analysis of statistical significance. **B.** Relative gene expression of FSP-1, Vimentin and Collagen I by qPCR in the scrambled control, scrambled transfected TGFβ1-stimulated (10 ng/ml) and TGFβ1-stimulated with (10 ng/ml) Angplt4 or Angptl3 siRNA-transfected cultured HK-2 cells. Data are mean ± SEM. One-way Anova with Tukey’s multiple comparison post hoc test was used to calculate statistical significance. **C.** Analysis of human ANGPTL4 expression in different tissues using Genotype-Tissue Expression (GTEx) database. TPM (transcript per million reads) is shown.

**Figure S5**

Masson Trichrome (MTS), Sirius red and PAS staining in kidneys of nondiabetic control and nondiabetic Angptl4^-/-^ mice. Representative histologic images are shown (original magnification 200X). Relative area of fibrosis (%), relative collagen deposition (%) and glomerular surface area were measured using ImageJ. Scale bar 100 µm. n=5/group. Data are mean ± SEM. Student t-test (unpaired two-tailed) was used for analysis of statistical significance.

**Figure S6**

A. In the UUO model, left kidneys were ligated in control littermates and Angptl4 mutant mice. Two days after UUO, four consecutive, low-dose (50 mg/kg ip) STZ injections were given. On day 11 kidneys were excised. Masson trichrome staining (MTS) in the contralateral non-operated kidney and UUO-operated kidneys in control and Angptl4 mutant mice were analyzed. Representative histologic images are shown; (original magnification 300X). Area of fibrosis (%) was measured using ImageJ program. n=6 mice/group combined from two independent experiments. Scale bar 100 µm. **B.** Blood glucose and kidney weight/body weight measurement in the indicated groups. Data are mean ± SEM. One-way Anova with Tukey’s multiple comparison post hoc test was used to calculate statistical significance. Significance-*p<0.05. **C.** Basal oxygen consumption rates (OCR) in kidneys from non-diabetic and diabetic wild-type and Angptl4^-/-^ mice were measured using the Seahorse Flux Analyzer (Agilent) according to the manufacturer’s instructions. Respiration rates were measured three times using an instrument protocol of 3-minute mix, 2-minute wait, and 3-minute measure. Flux rates were normalized to tissue weight. Experiments were repeated in 3 mice/condition. Data are mean ± SEM. One-way Anova with Tukey’s multiple comparison post hoc test was used to calculate statistical significance. Significance-*p<0.05.

**Figure S7**

A. Relative gene expression analysis of *Angptl4* in isolated podocytes from control and pmut mice. Data are mean ± SEM. n=7/group. Relative gene expression analysis of *Angptl4* in isolated tubules of control and tmut mice. Data are mean ± SEM. n=6/ group. Student t-test (unpaired two-tailed) was used for analysis of statistical significance. Significance *p<0.05. **B**. Graphical representation of coculture experiments in endothelial cells that were co-cultured with podocytes isolated from diabetic control and diabetic pmut mice. **C.** qPCR gene expression analysis of *αSMA*, *FSP-1* and *TGFβR1* in endothelial cells which were cocultured with podocytes from control mice and pmut mice**. D.** Measurement of IL-1β, IL-6, IL-10, IL-17 and TNFα levels in the cell supernatant. **E.** Radiolabeled [^14^C]palmitate uptake and oxidation, were measured in indicated groups. CPM of each sample was counted. **F.** ATP measurement in the endothelial cells which were co-cultured with podocytes either from diabetic control or diabetic pmut mice. **G.** Endothelial cell permeability assay in the indicated groups was analyzed by measuring FITC-dextran**. H.** qPCR gene expression analysis of miR-29a-3p, miR-29b-3p and miR-29c-3p in endothelial cells which were cocultured with podocytes from diabetic control and diabetic pmut mice. Data are mean ± SEM. Student t-test (unpaired two-tailed) was used for analysis of statistical significance. Significance- **p*<0.05.

**Figure S8**

**A**. Graphical representation of co-culture experiments in endothelial cells which were co-cultured with tubules isolated from diabetic control or diabetic tmut mice. **B**. qPCR gene expression analysis of *αSMA*, *FSP-1* and *TGFβR1* in endothelial cells which were co-cultured with tubules from control mice and tmut mice. **C**. Measurement of IL-1β, IL-6, IL-10, IL-17 and TNFα levels in the cell supernatant. **D.** Radiolabeled [^14^C]palmitate uptake and oxidation, were measured in indicated groups. CPM of each sample was counted. **E**. ATP measurement in endothelial cells which were co-cultured with tubules either from diabetic control or diabetic tmut mice. **F.** Endothelial permeability assay in the indicated groups was analyzed by measuring the concentration of FITC-dextran. **G**. qPCR gene expression analysis of miR-29a-3p, miR-29b-3p and miR-29c-3p in endothelial cells which were co-cultured with tubules from diabetic control and diabetic tmut mice. Data are mean ± SEM. Student t-test (unpaired two-tailed) was used for analysis of statistical significance. Significance- **p*<0.05.

**Figure S9**

A. *Angptl4* mRNA expression in kidneys from linagliptin-treated diabetic mice and untreated diabetic mice. n=7/group. **B.** *Angptl4* mRNA expression in kidneys of neutralizing integrin β1 antibody (N-integrin β1)-injected diabetic mice and IgG- injected diabetic mice. n=5/group. Data are mean ± SEM. Student t-test (unpaired two-tailed) was used for analysis of statistical significance. Significance-*p<0.05.

**Figure S10**

**A.** Physiological characteristics including blood glucose, kidney weight/body weight and albumin-to-creatine ratio were analyzed in control LNA- and miR-29 LNA-treated nondiabetic and diabetic mice. n=7 in the nondiabetic group and n=6 in the diabetic group. **B.** Relative gene expression analysis of miR-29a-3p, miR-29b-3p, and miR-29c-3p in the kidneys of control LNA- and miR-29 LNA-treated nondiabetic and diabetic mice. Data are mean ± SEM. n=7 in the nondiabetic group and n=6 in the diabetic group. One-way Anova with Tukey’s multiple comparison post hoc test was used to calculate statistical significance. Significance-*p<0.05.

**Figure S11**

A. Physiological characteristics including body weight, blood glucose, kidney weight/body weight, albumin-to-creatine ratio and mean blood pressure were analyzed in control, diabetic and N-acetyl-D-mannosamine-treated diabetic mice. n=6/group. **B.** Representative histologic images from MTS, Sirius red and PAS staining in kidneys of control, diabetic and N-acetyl-D-mannosamine-treated diabetic mice (original magnification 300X). Relative area of fibrosis (%), relative collagen deposition(%) and glomerular surface area were measured using ImageJ. Scale bar 70 µm. Data are mean ± SEM. One-way Anova with Tukey’s multiple comparison post hoc test was used to calculate statistical significance. Significance-*p<0.05.

**Figure S12**

**A.** Representative histologic images from Sirius red staining in kidneys from control, angiotensin-converting enzyme inhibitor imidapril (ACEi)-treated, N-seryl-acetyl-lysyl-proline (AcSDKP)-treated, or combined ACEi+AcSDKP-treated diabetic mice. Associated relative collagen deposition (%) is shown. Scale bar 50 µm. **B.** Relative gene expression levels of Angptl4 in the indicated groups. Data are mean ± SEM. One-way Anova with Tukey’s multiple comparison post hoc test was used to calculate statistical significance. Significance-*p<0.05.
