## Supplementary figures and images for "Renal Angptl4 is a key fibrogenic molecule in progressive diabetic kidney disease"

### Figure S1

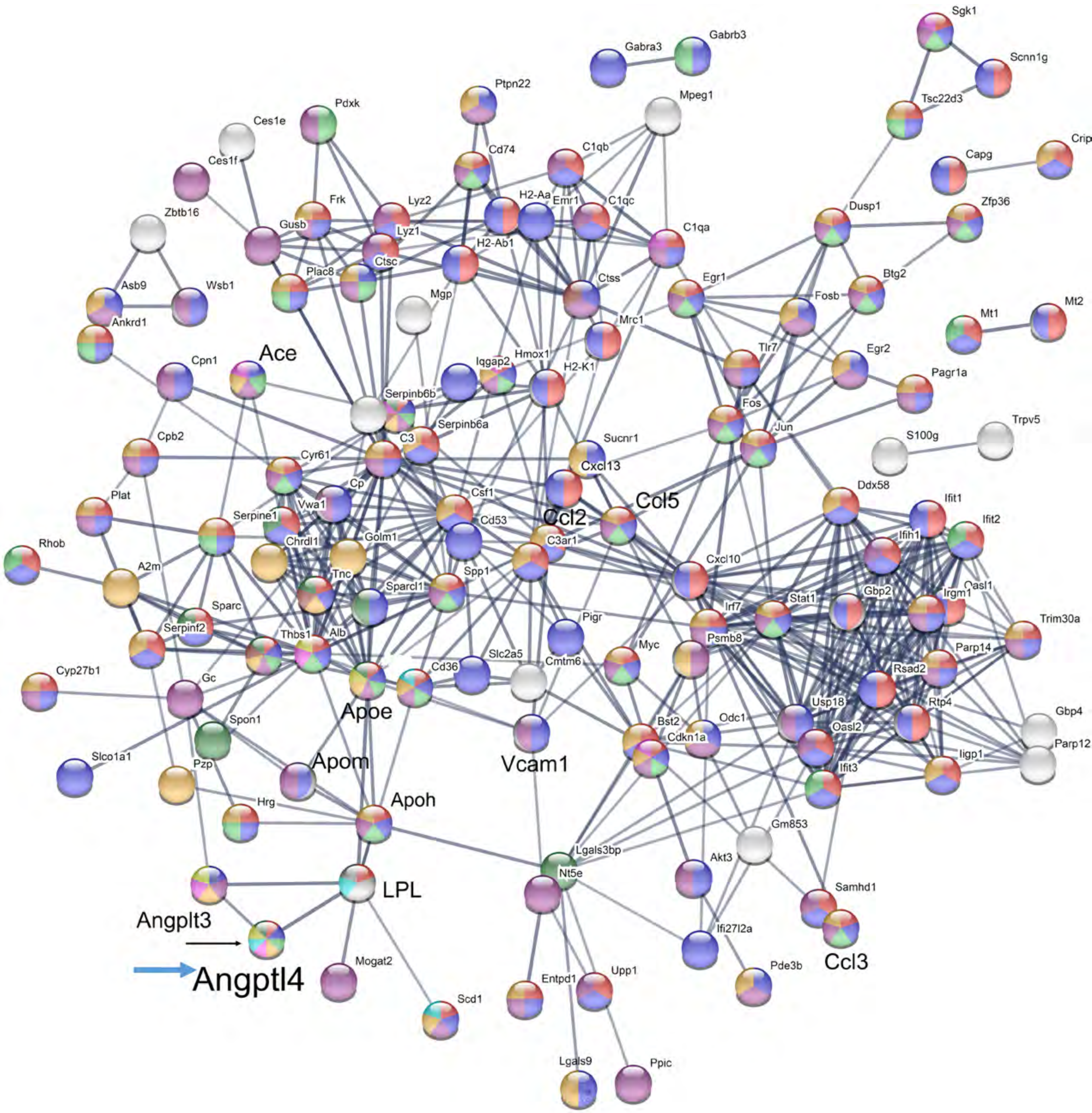

### Figure S2

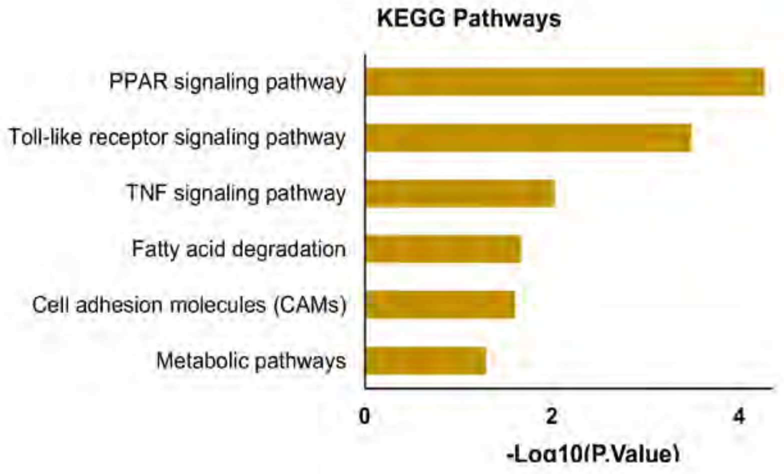

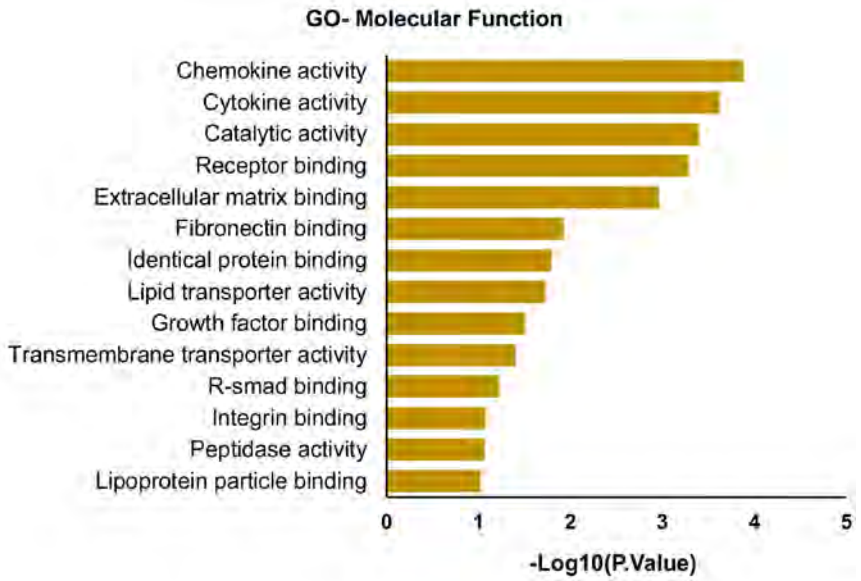

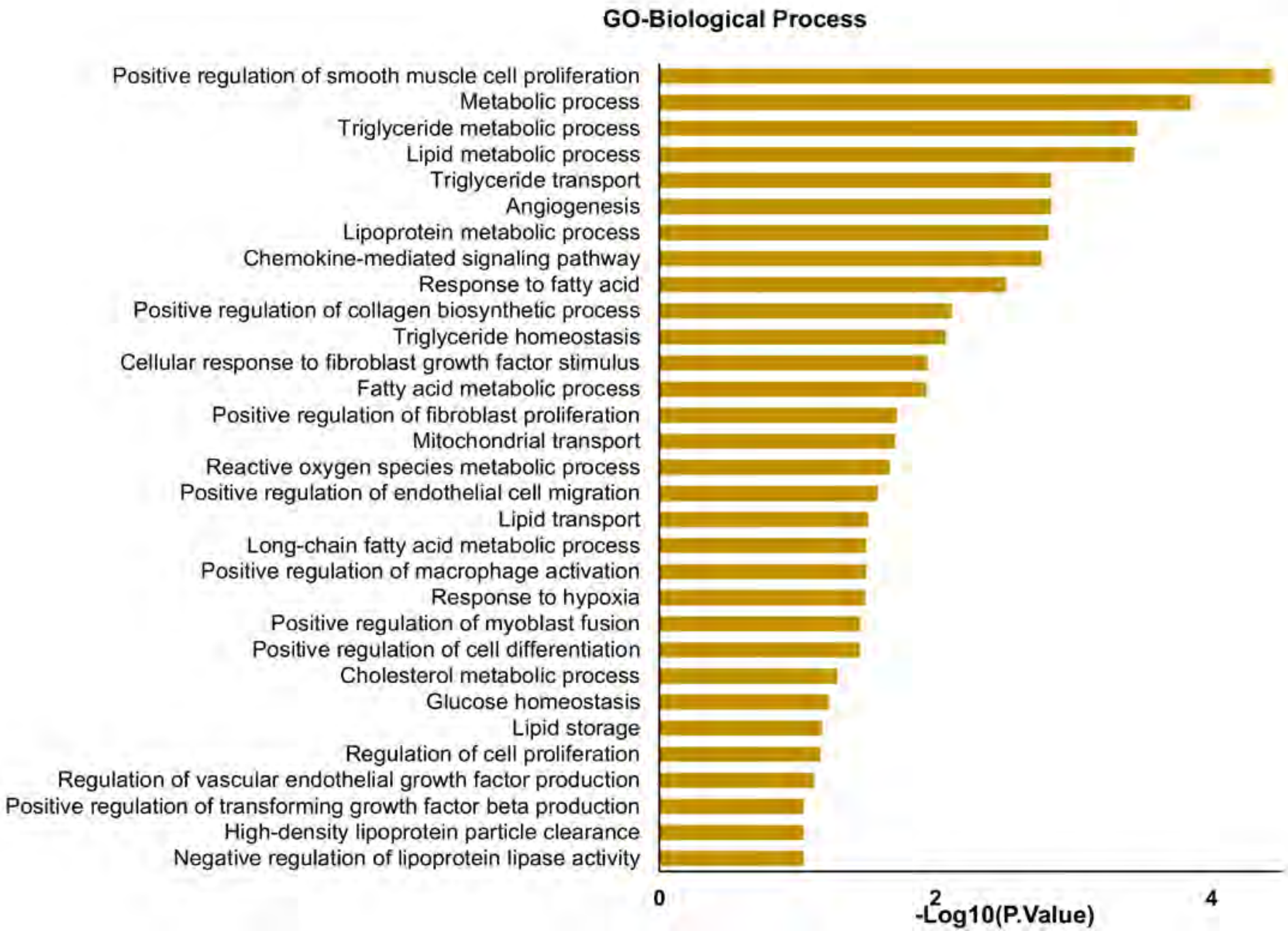

### Figure S3

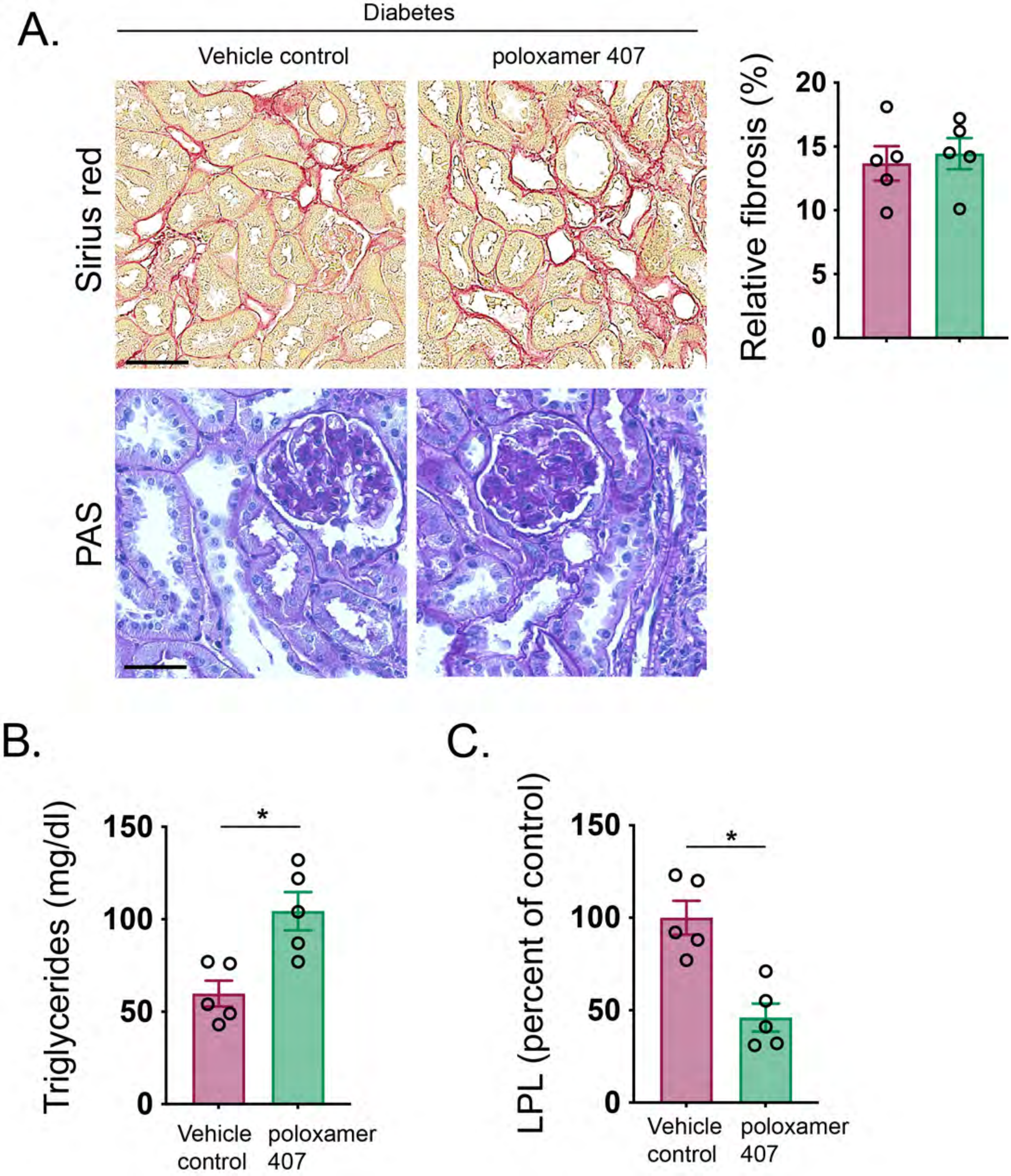

### Figure S4

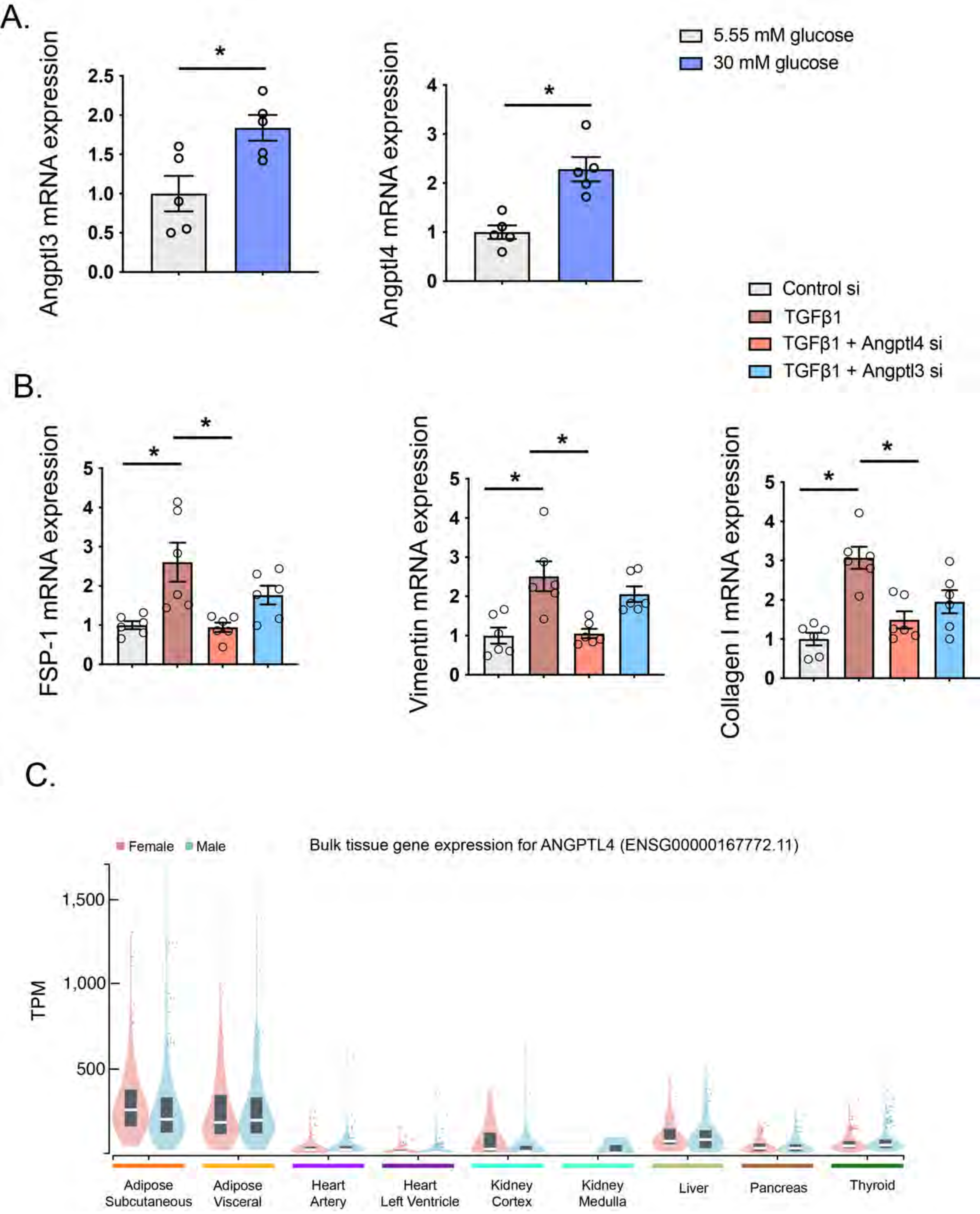

### Figure S5

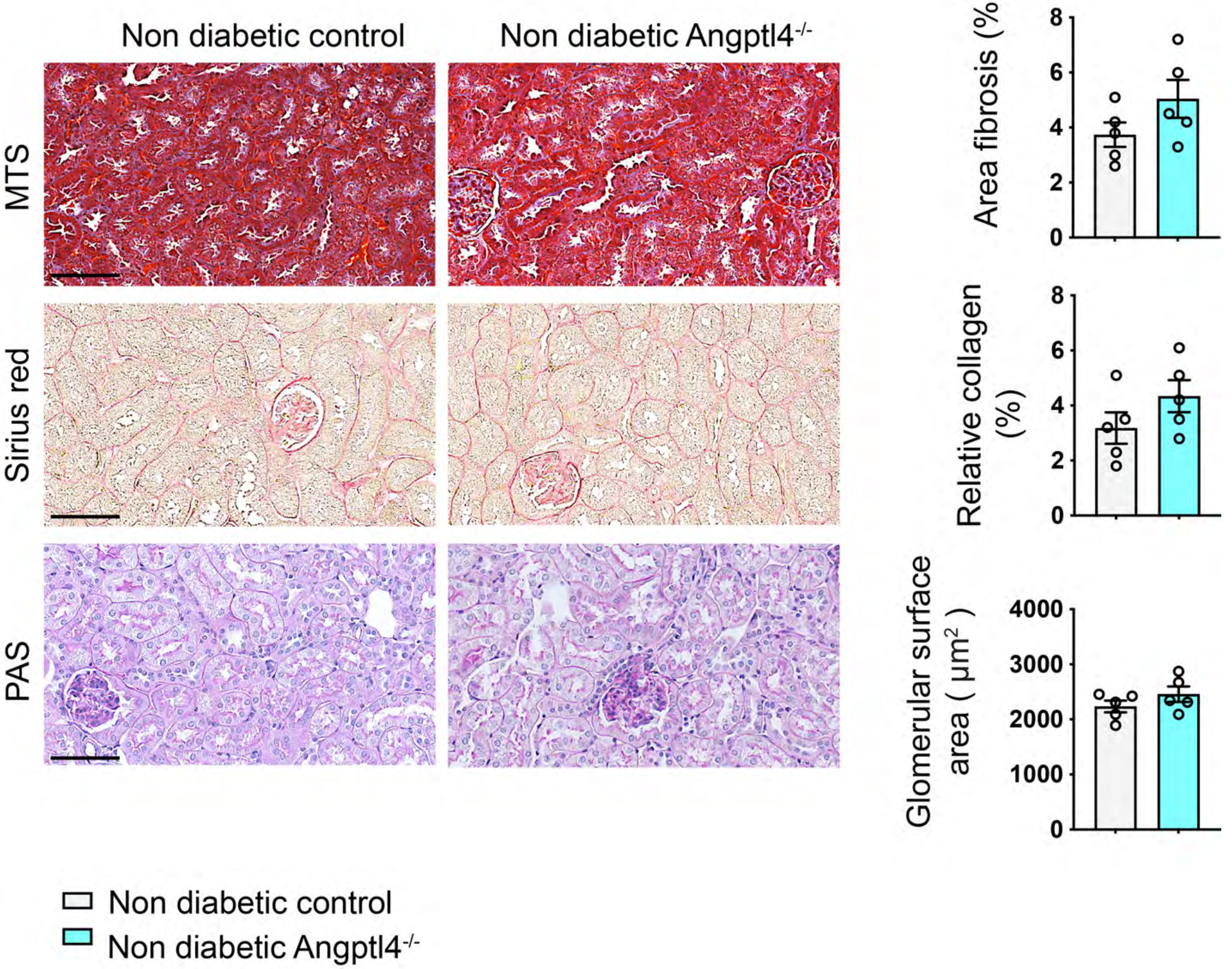

### Figure S6

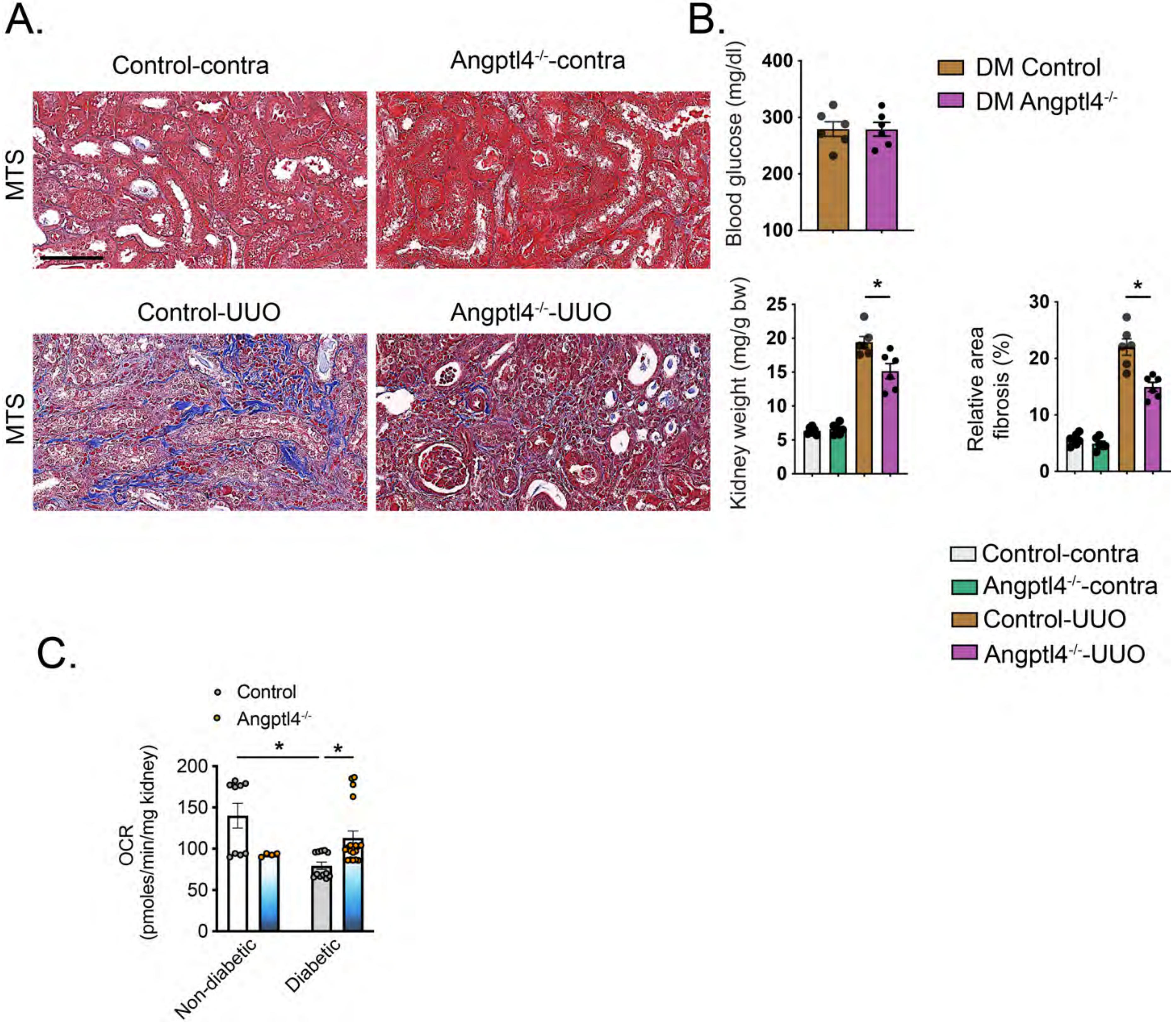

### Figure S7

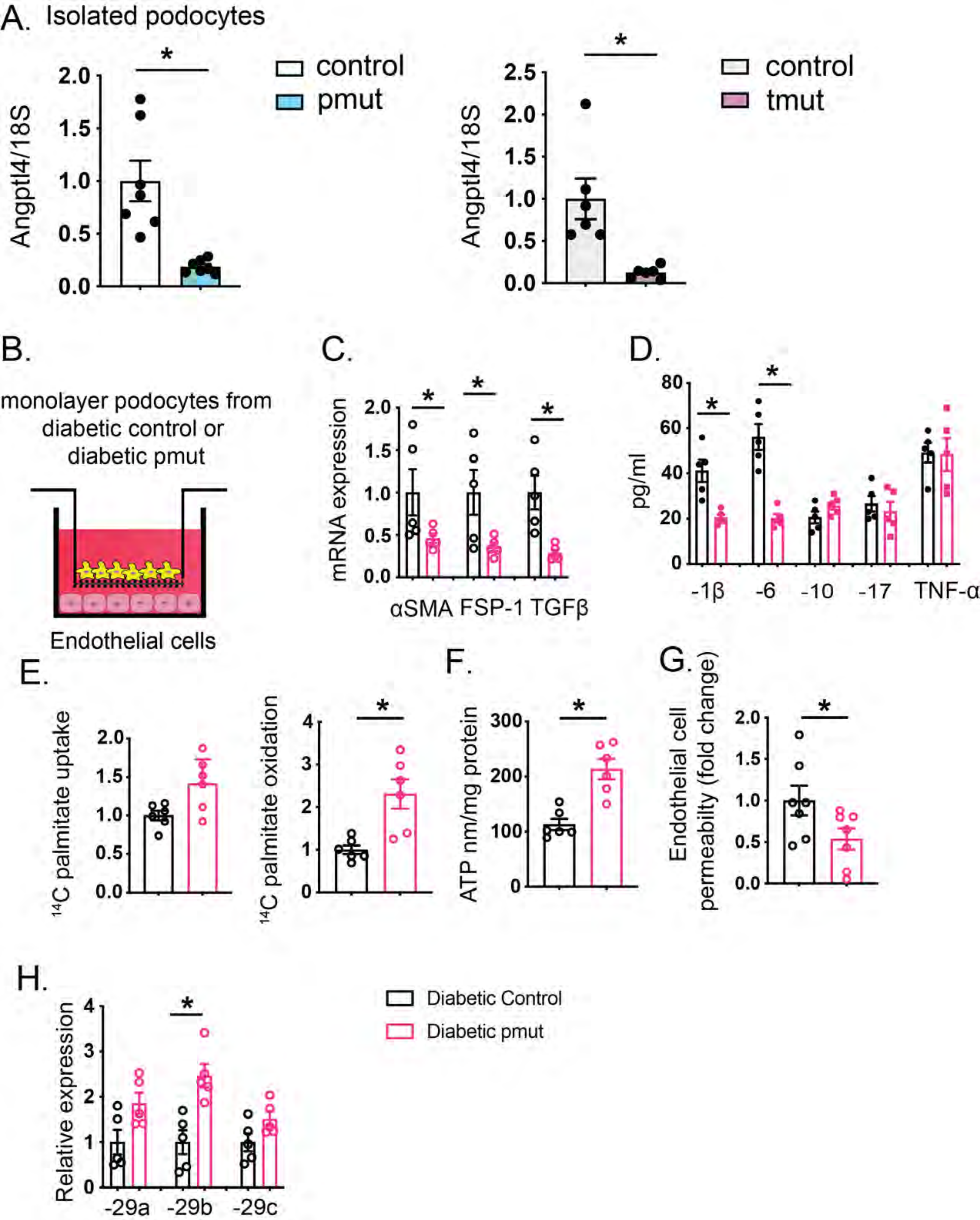

### Figure S8

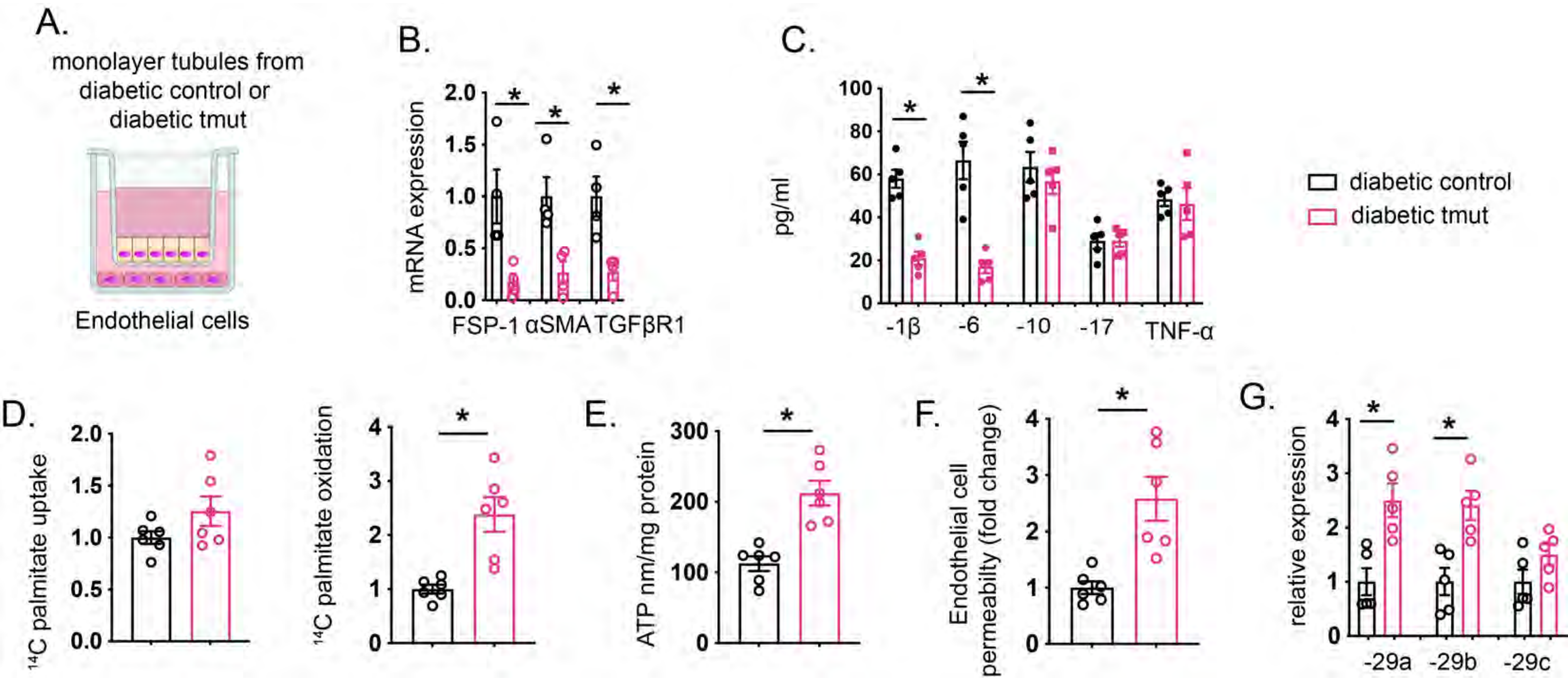

### Figure S9

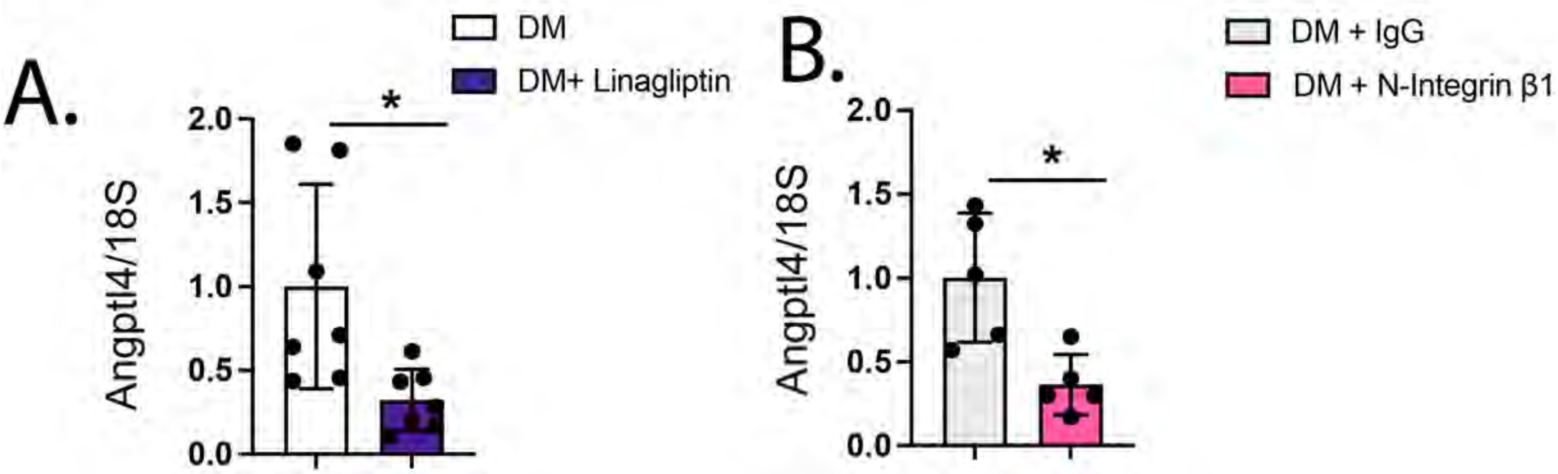

### Figure S10

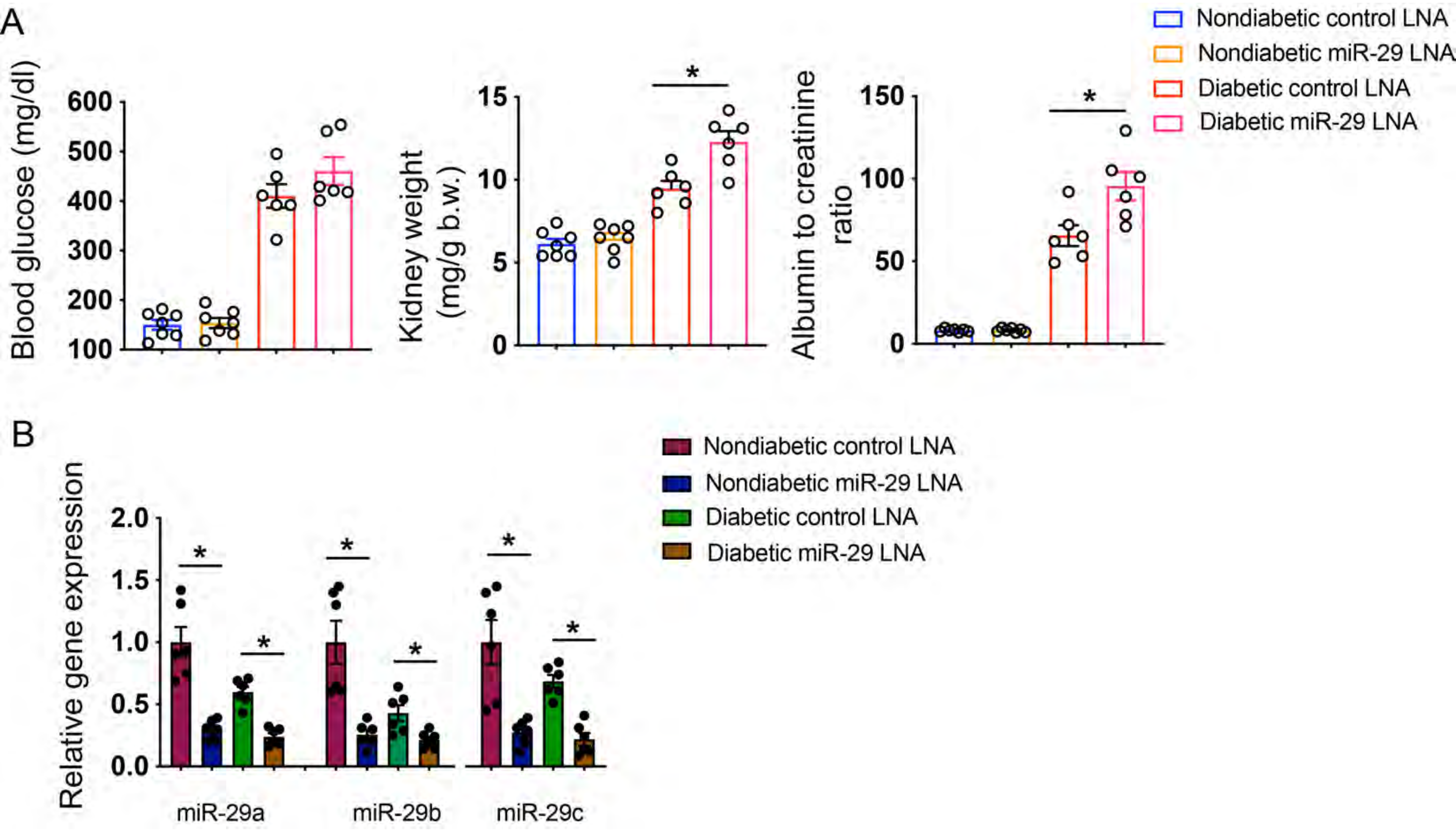

### Figure S11

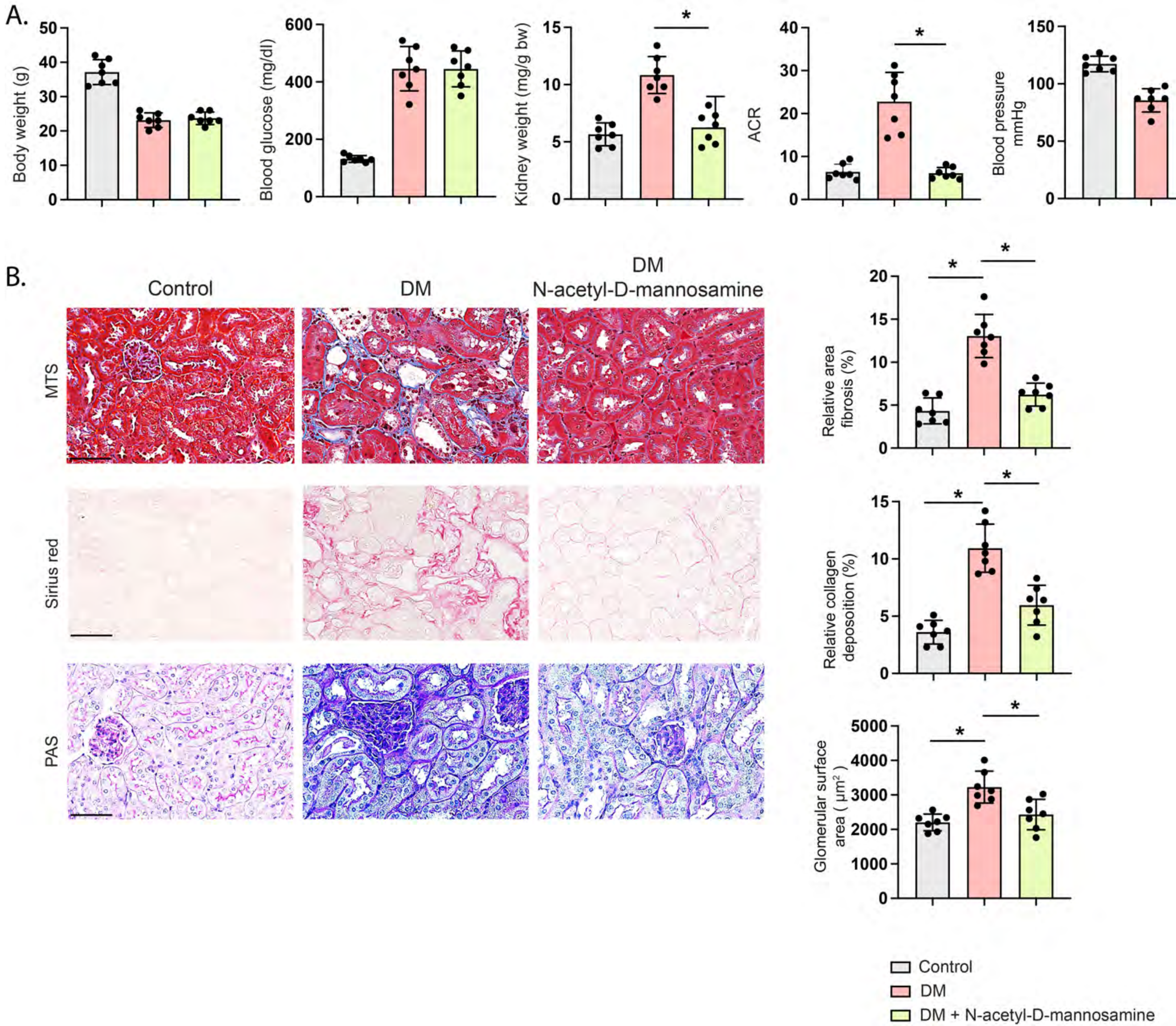

### Figure S12

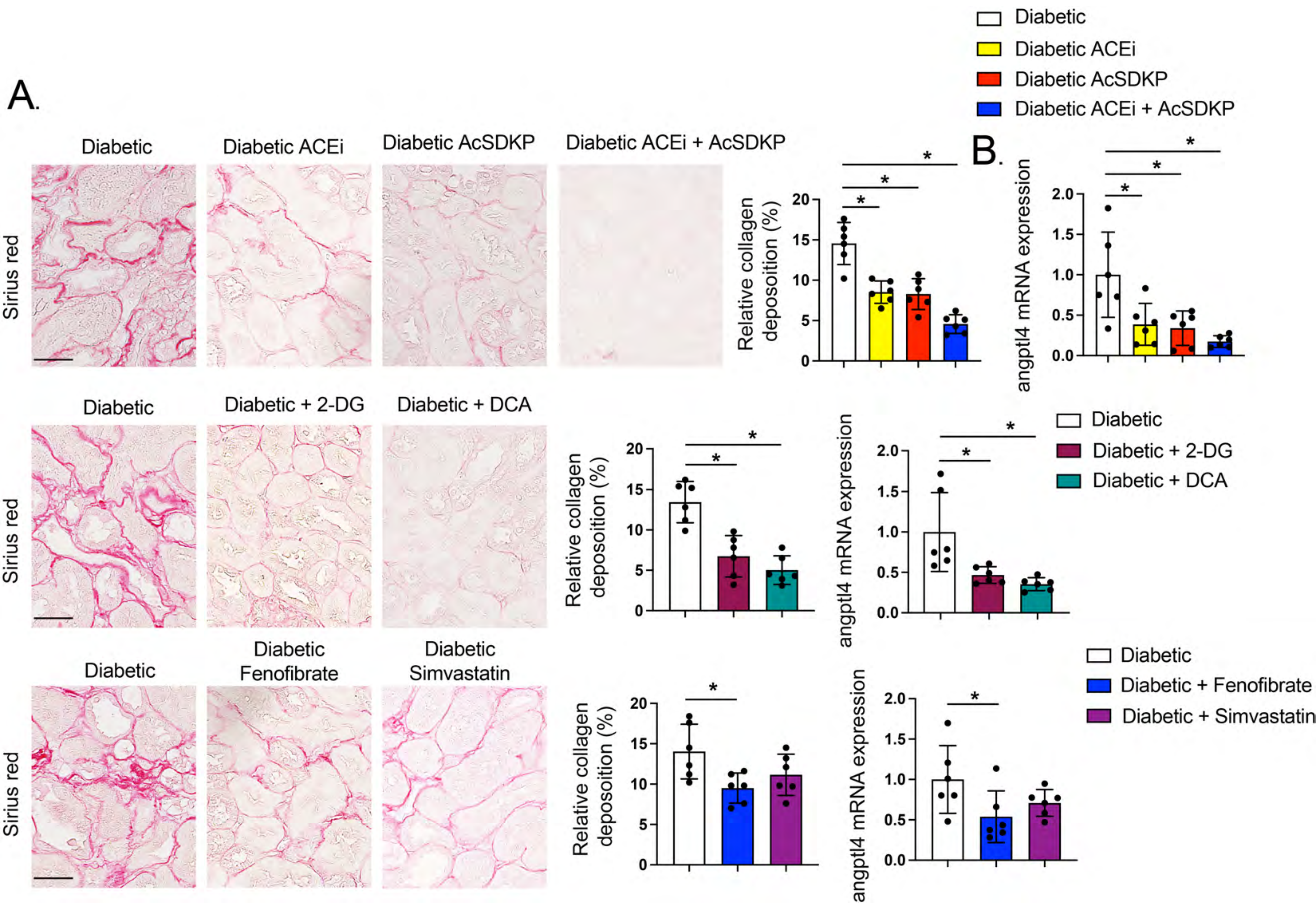
